# Glycine Rich Segments Adopt Polyproline II Helices Which May Contribute to Biomolecular Condensate Formation

**DOI:** 10.1101/2020.07.30.229062

**Authors:** Miguel Mompeán, Bethan S. McAvan, Sara S. Félix, Miguel Treviño, Javier Oroz, David Pantoja-Uceda, Eurico J. Cabrita, Andrew J. Doig, Douglas V. Laurents

## Abstract

Many intrinsically disordered proteins contain Gly-rich regions which are generally assumed to be disordered. Such regions often form biomolecular condensates which play essential roles in organizing cellular processes. However, the bases of their formation and stability are still not completely understood. Considering NMR studies of the Gly-rich *H. harveyi* “snow flea” antifreeze protein, we recently proposed that Gly-rich sequences, such as the third “RGG” region of Fused in Sarcoma (FUS) protein, may adopt polyproline II helices whose association might stabilize condensates. Here, this hypothesis is tested with a polypeptide corresponding to the third RGG region of FUS. NMR spectroscopy and molecular dynamics simulations suggest that significant populations of polyproline II helix are present. These findings are corroborated in a model peptide Ac-RGGYGGRGGWGGRGGY-NH_2_, where a peak characteristic of polyproline II helix is observed using CD spectroscopy. Its intensity suggests a polyproline II population of 40%. This result is supported by data from FTIR and NMR spectroscopies. In the latter, NOE correlations are observed between the Tyr and Arg, and Arg and Trp side chain hydrogens, confirming that side chains spaced three residues apart are close in space. Taken together, the data are consistent with a polyproline II helix, which is bent to optimize interactions between guanidinium and aromatic moieties, in equilibrium with a statistical coil ensemble. In cells, the polyproline II population of these peptides could be augmented by binding profilin protein or SH3, WW or OCRE domains, association with RNA or assembly into polyproline II helical bundles. These results lend credence to the hypothesis that Gly-rich segments of disordered proteins may form polyproline II helices which help stabilize biomolecular condensates.

## Introduction

Biomolecular condensates, also known as membraneless organelles, comprise a diverse family of cellular structures performing many essential functions (Banani *et al.*, 2017), mainly related to RNA regulation. Compared to classic organelles like the Golgi body or the lysosome, membrane-less organelles are much more dynamic (Boeynaems *et al.*, 2018). They can quickly form or dissociate within minutes in response to changing cellular conditions such as stress. Such agile assembly and disassembly is possible because these condensates form by liquid-liquid phase separation (LLPS). This process excludes most water molecules and is driven by more favorable interactions within the condensate relative to those in bulk solution. While individually weak and transient, these interactions are collectively strong enough to drive LLPS.

Over the last decade, many seminal studies have uncovered and characterized the contributions of several classes of weak interactions (Wang *et al.*, 2018) to condensate formation. Some, such as polyproline II helices + SH3 domains (Li *et al.*, 2012 & Amaya *et al.*, 2018), the N-terminal domain of TDP-43 (Mompeán *et al.*, 2017; Wang *et al.*, 2018), and α-helix + α-helix associations, operate on the level of protein domains or secondary structure. Other favorable attractions arise between small groups of atoms; namely: cation-π, π-π (Vernon *et al.*, 2018), cation-anion, and hydrophobic interactions. Interactions between protein and RNA by charge-charge and cation-anion interactions also modulate the formation of most membraneless organelles.

Despite these key advances, our understanding of the factors driving membraneless organelle formation is incomplete and the brisk pace of discovery in this field suggests that additional types of interactions are yet to be characterized. Recent work has shown the key role of RGG segments in promoting condensate formation by interacting with S/G,Y,S/G segments in Fused in Sarcoma (FUS) and related proteins (Wang *et al.*, 2018). In fact, the presence of the RGG segments lowered the critical concentration for condensate formation by a factor of approximately twenty. In addition to FUS, a number of proteins sharing some activities and most features of its domain organization, such as transactive DNA-binding response protein of 43 kDa (TDP-43), also contain glycine-rich intrinsically disordered domains. Whereas glycine is long known to destabilize α-helices and β-strands, it is essential to the structure and stability of the polyproline II triple helix of collagen, as well as polyproline II helical bundles, such as the snow flea antifreeze protein (Pentelute *et al.*, 2008). The small size of glycine is key to polyproline II inter-chain helix packing in these structures. Composed of sequence motifs Xaa-Gly-Gly, central helices in polyproline II helical bundles (Pentelute *et al.*, 2008; Singh, *et al.*, 2017; Warkentin *et al.*, 2017) require two consecutive glycines as each polyproline II helix is closely packed and hydrogen bonded to up to four others.

The folding of glycine-rich polypeptides is strongly opposed by loss of conformational entropy, considering that glycine occupies broad regions of the Ramachandran map when disordered but is confined to a tiny zone of φ,ψ space in the polyproline II conformation (Gates *et al.*, 2017). Recently, spectroscopic and computational evidence of glycine residues in polyproline II helical bundles forming weak Cα-H---:O=C hydrogen bonds were reported (Gates *et al.*, 2017; Treviño *et al.* 2018). Such interactions could add up to a substantial stabilizing contribution, which could compensate for the entropic cost of folding. Sequence analysis revealed that many proteins, such as FUS or nucleolin, which are assumed to be disordered, have Xaa-Gly-Gly motifs. We proposed that they may form polyproline II helices (Treviño *et al.* 2018). The aim of the study reported here is to test this hypothesis using model peptides based on the third RGG segment of FUS. Our experimental characterization, via circular dichroism (CD), Fourier transform infrared (FTIR) and nuclear magnetic resonance (NMR) spectroscopies, evinces that this peptide can adopt a polyproline II helix.

## Results

### NMR spectroscopic

characterization of the peptide RGG3FUS, whose sequence is RGGRGGYDRGGYRGGRGGDRGNYRGGRGGD, lead to the successful assignment of ^1^H and ^13^C nuclei at different pH values; namely, 2.53, 7.25, and 9.96, which are listed in **Tables S1-S3**. These three pH values were chosen to test if the charged state of Asp and Tyr residues affect the formation of ordered structure. The assigned 2D ^1^H-^13^C HSQC spectrum acquired at pH 9.96 is shown in **Sup. Fig. 1** and the ^1^HN-^1^Hα zone of the 2D ^1^H-^1^H COSY, TOCSY and NOESY spectra at pH 2.53 and 7.25 values are shown in **Sup. Fig. 2**. Spectra at pH 9.96 are not shown because the HN signals disappear due to exchange with water under alkaline conditions. Relatively small ^1^Hα and ^13^Cα conformational chemical shifts are observed at acidic, neutral and alkaline pH; this means there is no significant population of α-helix or β-strand, and points to isolated polyproline II helix or disordered conformers (**Fig. 1A**). Moreover, ^1^HN-^1^Ht coupling constants (^3^J_HNHA_), which provide an orthogonal measure of secondary structure, are consistent with φ angle values typical of extended backbone conformations, such as β-strands or polyproline II helices (**Fig. 1B**, Pardi *et al.*, 1984). These two results together suggest a conformational ensemble with polyproline II helices over a broad range of pH conditions, from acidic to alkaline.

**Figure 1:**
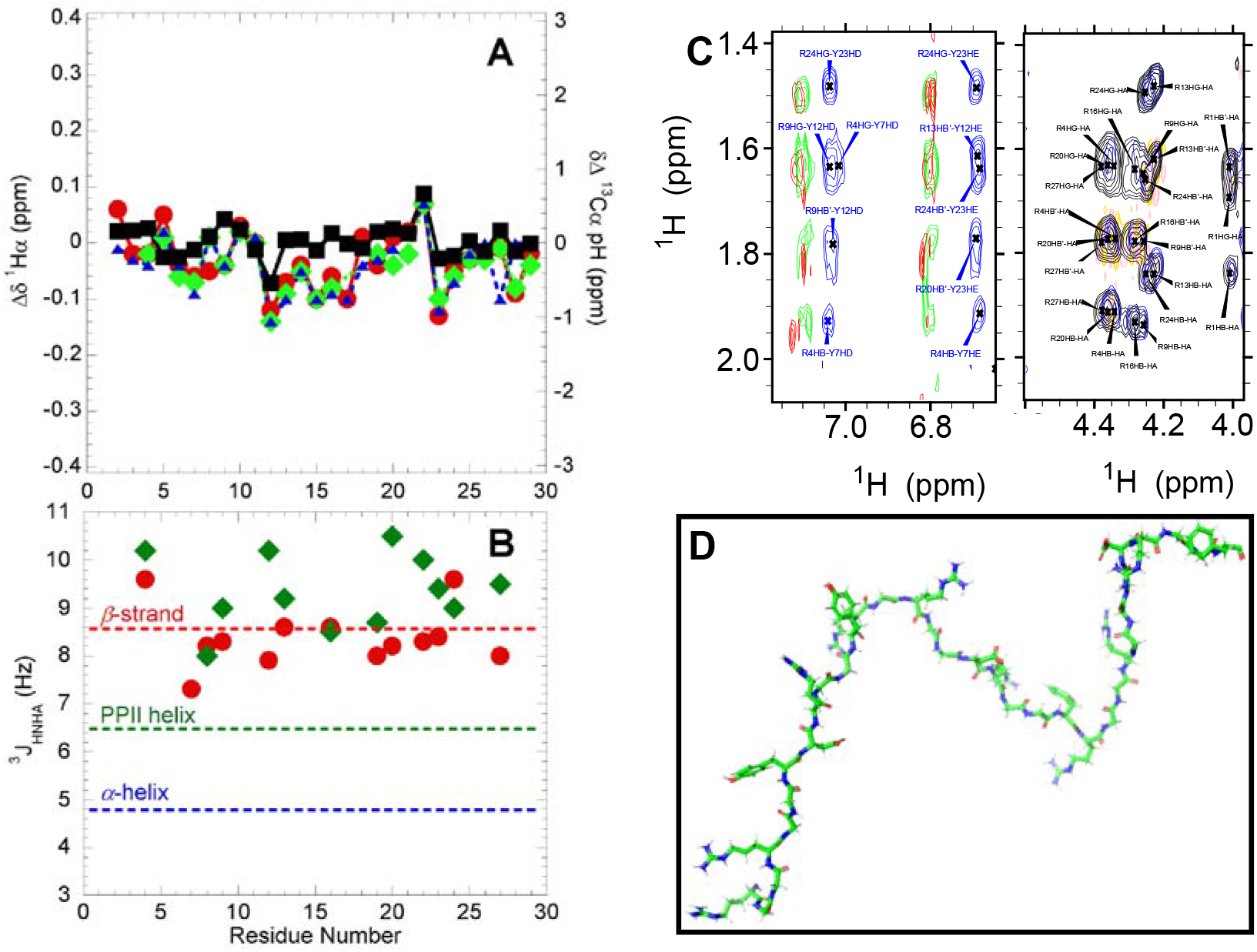
NMR Characterization of RGG3FUS. **A.** Conformational ^1^Hα chemical shifts (Δδ ^1^Hα, left y-axis) for RGG3FUS measured at 5 °C and pH 2.53 (**red** circles), pH 7.25 (**green** diamonds) and pH 9.96 (**blue** triangles). Conformational ^13^Cα chemical shifts (Δδ ^13^Cα, right y-axis) measured at 5 °C and pH 9.96 (**black** squares). The y-axis scales (0.82 ppm for ^1^Hx and 6.2 ppm for ^13^Cα) cover the full ranges expected for 100% α-helix (−0.41 ppm for ^1^Hx, +3.1 for ^13^C) to 100% β-strand (+0.41 for ^1^Hx, −3.1 for ^13^Cα). N- and C-terminal residues, which are perturbed by end effects, are not plotted. **B.** ^1^HN-^1^H. coupling constants for RGG3FUS at pH 2.53 (**red** circles) and pH 7.25 (**green** diamonds). The mean values for α-helix, PPII helix and β-strand are marked by **blue**, **green** and **red** dashed lines, respectively. **C.** *Left panel:* 2D ^1^H-^1^H NOESY spectra of RGG3FUS recorded at 5°C and pH 2.53 (**red**), 7.25 (**green**) and 9.96 (**blue**). Crosspeaks arising from ^1^H contacts between Arg and Tyr side chains are labeled. *Right panel:* Arg side chain assignments: COSY correlations are colored **gold**(+) and **pink**(−), TOCSY=**black**, NOESY=**blue**. **D.** Representative conformers of RGG3FUS. Some polyproline II helical turns, stabilized by Arg-Tyr and Arg-Asp side chain interactions, which are spaced by kinks, can be observed. At the N-terminus (lower left corner) a possible cation-π interaction between two Arg guanidinium moieties, disposed perpendicularly, may be present.

In general, side chain chemical shifts are close to their expected values for being exposed to solvent, suggesting there no stable hydrophobic core is formed. However, the side chain ^1^Hβ, ^1^Hγ and ^1^Hδ resonances of Arg 13 and Arg 24 are shifted upfield, which can be attributed to contacts with the aromatic rings of Tyr 12 and Tyr 23, respectively. Contacts between Arg and Tyr side chain protons are corroborated by the observation of NOE crosspeaks (**Fig. 1C**). Whereas the similar chemical shift values make it difficult to unambiguously assign these crosspeaks, some of them arise from non-sequential contacts and could be due to Arg and Tyr residues spaced *i, i+3* along the sequence, which would be consistent with a turn of polyproline II helix. Conformers of RGG3FUS containing these short polyproline II helices stabilized by side chain interactions, generated with CYANA, are shown in **Fig. 1D**. These structures are minor species in the conformational ensemble of the third RGG region of FUS.

### Molecular Dynamics simulations on RGG3FUS and design of a mimicking system

To further explore the ability of RGG sequences to sample PPII conformations, the structural models generated using CYANA, based on the above NMR insights, were submitted to three independent Molecular Dynamics simulations. The starting structures were let to evolve through distinct trajectories by setting different initial velocities, for total sampling time of 217 ns. As illustrated in **Figure 2**, PPII conformations are sampled by the various amino acid residues, which lends confidence to the hypothesis that PPII species may be populated in the third RGG of FUS (RGG3FUS).

**Figure 2.**
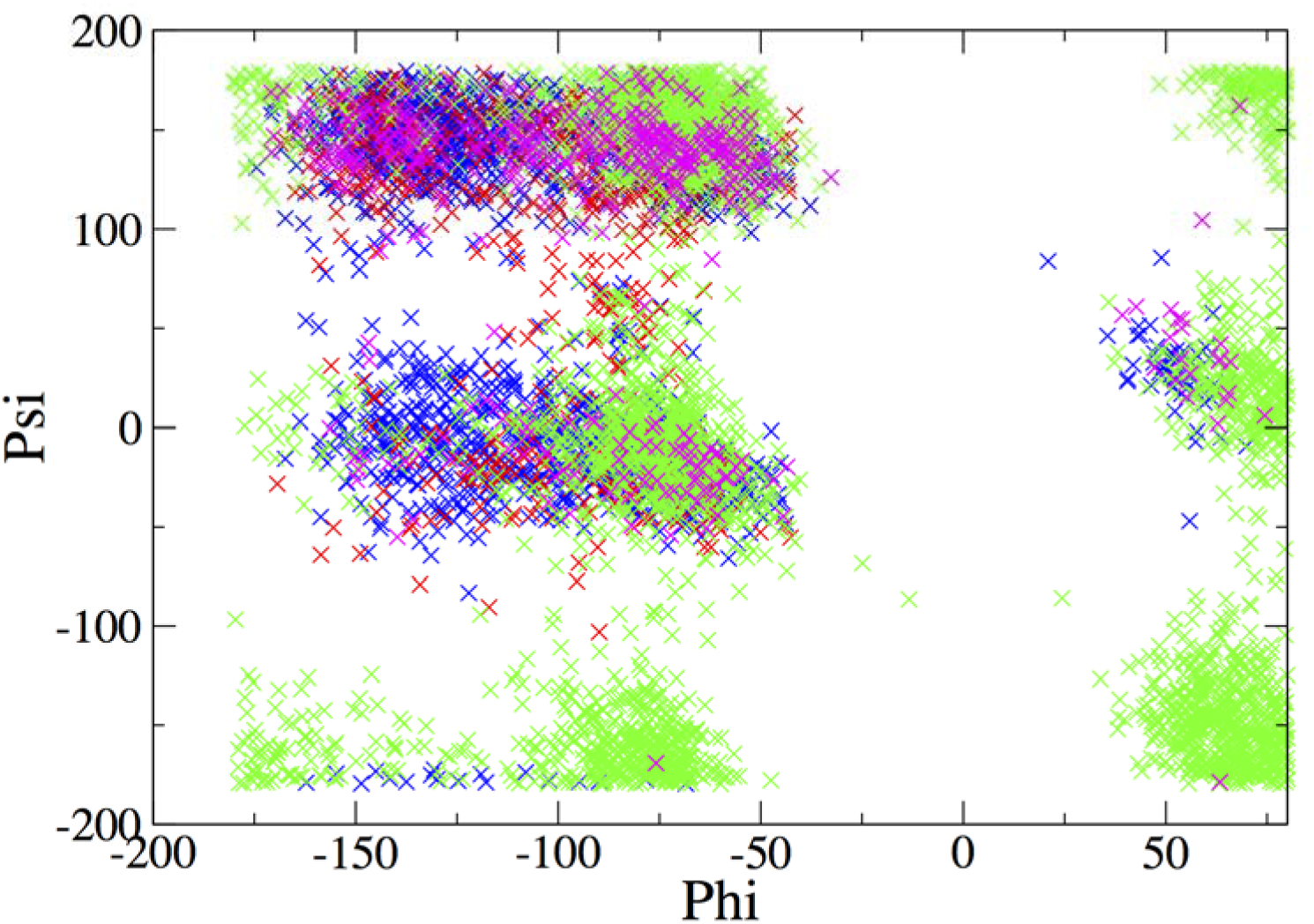
Molecular Dynamics simulations of RGG3 FUS. Ramachandran map showing sampling of the PPII region by Gly (**green**), Arg (**blue**), Asp (**red**) and Tyr (**magenta**) residues of RGG3FUS. The simulations were initiated using the NMR-driven model shown in **Fig. 1D**. **φ** and **ψ** angles are plotted from snapshots were retrieved every nanosecond from a total sampling time of 217 ns. The PPII region spans approximately −100°to −50° for **ϕ** and +110° to +190° for **ψ**.

The analysis of the third RGG motif of FUS in representative vertebrates indicates that the number of RGGY/F and RGGD tracts is conserved. Residues capable of forming attractive cation/**π** interactions spaced *i, i+3* are also observed in the C-terminal region of nucleolin, a nucleolar protein (Feric *et al.*, 2006; **Table 1**). Thus, based on the above NMR and modeling data, the sequence of RGG3FUS in *H. sapiens* and *G. Gallus*, R_473_GG<u>RGGY</u>DRGGY<u>RGGRGGD</u>RGNYRGG<u>RGGD</u>_506_, would afford such cation (*i*) / π (*i*+3) and cation (*i*) / anion (*i*+3) interactions (single and double underlined stretches, respectively). To test this hypothesis, we designed a mimicking system to scrutinize the presence of these interactions and the existence of PPII conformations. In particular, a peptide containing the four underscored segments in RGG3FUS was synthesized, with the following modification: a Tyr has been substituted by Trp at the center of the sequence to facilitate NMR analysis and obtain site specific assignments that are not possible for RGG3FUS. We call this system RGGmini, and its sequence is as follows: RGGYGGRGGWGGRGGY. At alkaline pH, *i* to *i+3* interactions between R and Y may form cation-anion contacts, whereas at any other pH values they could interact through *i* to *i+3* cation-ττ contacts.

**Table 1:**
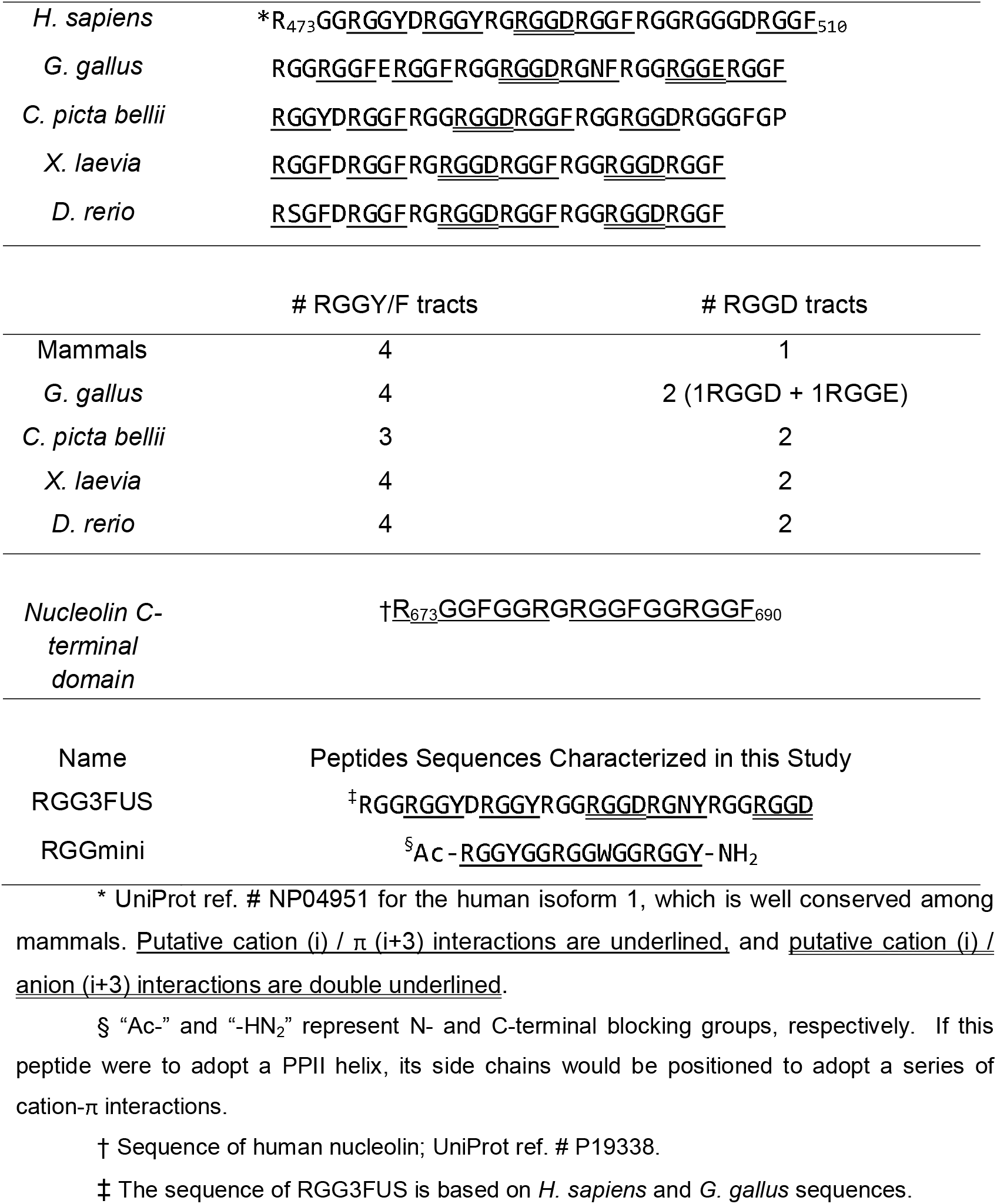
Sequence of the Third RGG Motif of FUS in Representative Vertebrates

### Circular dichroism spectra of RGG mini

The far UV CD spectra of RGGmini recorded at pH 10, 5°C revealed a positive band at 225 nm (**Figure 3A**), which is commonly observed for *bona fide* polyproline II helices (Tiffany & Krimm, 1968; Rucker & Creamer 2002; Lam & Hsu, 2003). No minima are detected at 218 or 222 nm which evidences a lack of β-strand or α-helical secondary structure. Similar spectra were observed when recorded at pH 3 or pH 8, except the shoulder from 240-250 nm less pronounced (data not shown). Based on the [θ] signal of +1000 deg·cm^2^·mol^−1^ at 222 nm, the population of polyproline II helix can be estimated to be about 40% using the formula reported by Stellwagan and co-workers (Park *et al.*, 1997), with 60% disordered conformers. This positive CD signal weakens at higher temperatures (**Figure 3A**). This is consistent with a loss of polyproline II helix due to thermal denaturation. However, aromatic moieties locked in a fixed structure can also produce positive CD bands (Chakrabartty *et al.*, 1994). To test for fixed aromatic side chains, we recorded near UV-CD spectra at 5°C and 60°C. No significant peaks were detected (**Figure 3B**). This suggests that the far UV-CD peak centered at 225 nm does arise from polyproline II helical structure. Nevertheless, it is possible that two fixed aromatic groups could produce peaks of opposite sign in the near UV CD spectra which cancel out, but sum to produce a positive peak in the far UV CD spectra. Thus, a possible contribution of fixed aromatic peaks could not be wholly disproved by the CD spectra.

**Figure 3.**
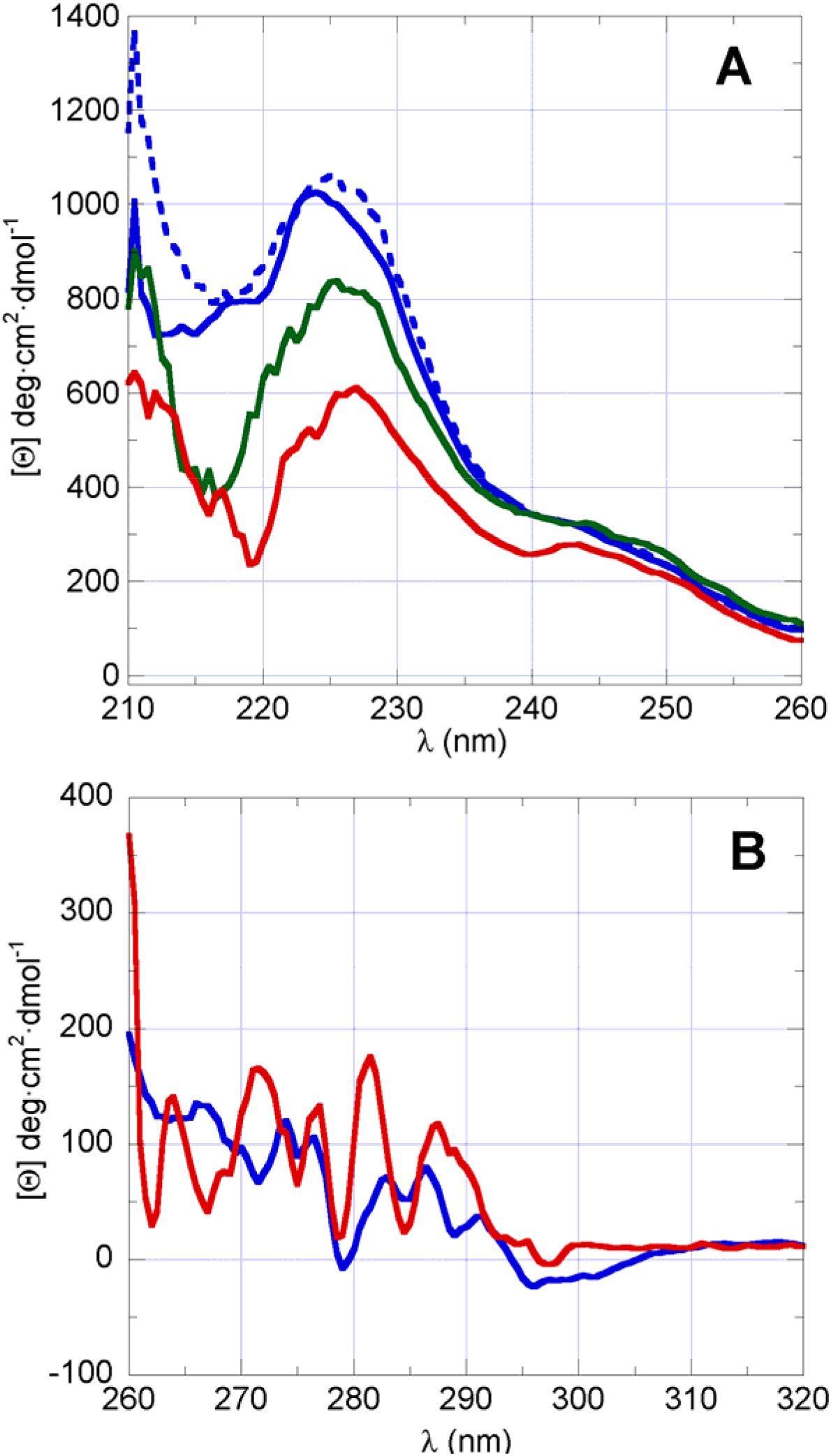
Circular Dichroism Characterization of RGGmini. **A**. Far UV-CD spectra of RGGmini at **0, 30, 60** °C (solid lines) and recooled to **0** °C (dashed line), pH 10. **B**. Near UV-CD spectra recorded at **5** and **60** °C, pH 10.

### Fourier Transform Infrared Spectroscopy

While CD and FTIR spectroscopies both provide information on protein secondary structure, FTIR produces spectral bands for the peptide free of contributions from aromatic groups. In other words, while the near-UV and far-UV regions in the CD spectra inform on aromatic and aromatics+backbone contributions, respectively, FTIR has the advantage that it reports solely on the backbone conformation. The analysis of FTIR spectra of RGGmini show evidence for polyproline II helix at pH 10, where *i* to *i+3* cation-anion RY interactions are inferred from our NMR-driven structural models and CD data, and at pH 3.5, where *i* to *i+3* R-Y contacts are cation-ττ (**Figure 4**). Although there have been relatively few studies of polyproline II helices by infrared spectroscopy, a recent study of the glassy hair protein KAP8.1, which is likely to adopt a bundle of four polyproline II helices, reported IR bands at 1670 cm^−1^ and 1656 cm^−1^ which could be assigned to antiparallel β-strand or coil or polyproline II helix (Singh *et al.* 2017). Taking into account the increased signal upon heating observed in **Figure 4** at 1655 cm^−1^, we assign that peak to statistical coil. Considering the decreased peak intensity of the 1672 cm^−1^ peak at 60 °C and the lack of CD or NMR evidence for β-strands described above, we assign the 1672 cm^−1^ band to polyproline II helix.

**Figure 4:**
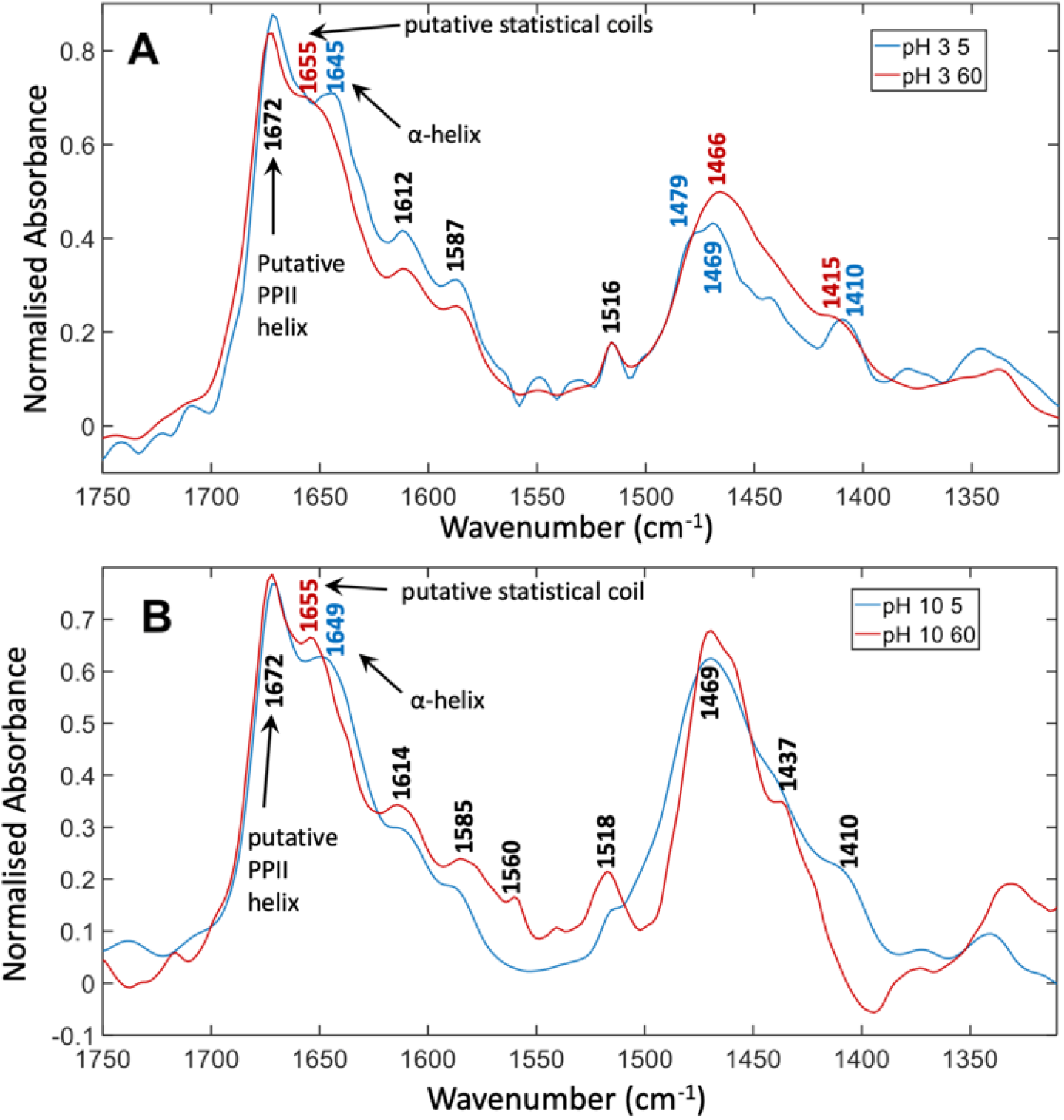
FTIR Spectral Characterization of RGGmini. **A**. FTIR spectra of 15 mM RGGmini at pH 3.5. **B**. FTIR spectra of 15 mM RGGmini at pH 10. The spectra shown are the average of three replicate spectra recorded with 256 scans each. Spectra recorded at **5°C** and **60°C** are colored **blue** and **red**; respectively.

### NMR Spectroscopy of RGGmini

To obtain atomic level conformational information, we characterized RGGmini by NMR spectroscopy. The assignments for the structurally relevant nuclei at pH 3 and pH 10 (**Tables S4 & S5, respectively**) were measured by recording and analysis of 2D ^1^H-^1^H COSY, TOCSY and NOESY spectra (**Figure 5A**) and 2D ^1^H-^13^C HSQC and ^1^H-^15^N HSQC spectra. Whereas broadened ^1^H-^15^N peaks pointed to the presence of dimeric sfAFP (Treviño *et al.*, 2018), here no signs of broadened peaks were observed for RGGmini. This suggests that RGGmini is monomeric as is consistent with its high net charge. The ^1^Hα conformational chemical shifts are small (**Figure 5B**); following the rationale of the NMR analysis for the parent RGG3FUS motif (**Figure 1**). This is consistent with the absence of α-helix, β-strands or polyproline II helical bundles, but would be expected for an isolated polyproline II helix or a statistical coil. Though the ^13^Cα conformational chemical shifts are also small, they are generally negative and are thus suggestive of partial **β**-strand or PPII helix formation (Treviño *et. al.*, 2018). Along this line, the ^1^HN-^1^Hx coupling constants (^3^J_HNHA_), obtained from the COSY spectrum are relatively large, which is consistent with β-strand or polyproline II helix, but not α-helix (**Figure 5C**, Pardi *et al.*, 1984). Taking together, these data evidence the presence of polyproline II helices within a conformational ensemble likely enriched in extended structures.

**Figure 5:**
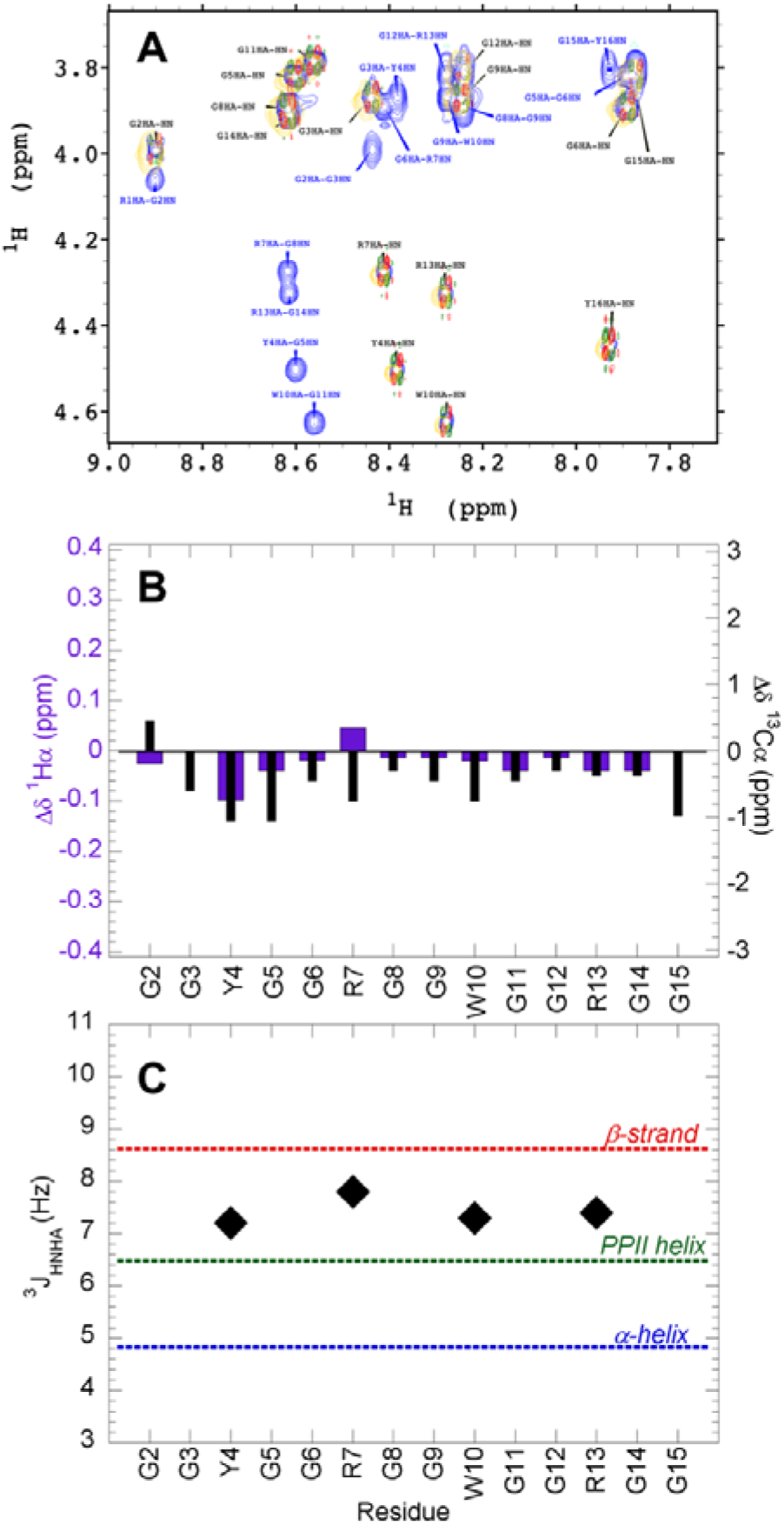
NMR Characterization of RGGmini (I) **A**. ^1^HN - ^1^H region of the 2D ^1^H-^1^H COSY (colored **green** and **red** for positive and negative correlations, respectively), TOCSY (**gold**) and NOESY (**blue**) spectra of RGGmini. Inter- and intra-residue correlations are colored **blue** and **black**, respectively. Multiplication factor for contours = (2)^1/2^, so every two steps is a factor of two. **B**. Conformational chemical shifts of ^1^H**α** (left y-axis, **purple** bars) and ^13^C**α** (right y-axis, **black** bars) of RGGmini. The y-axis scales (0.82 ppm for ^1^Ha and 6.2 ppm for ^13^C**α**) cover the full ranges expected for 100% -helix (−0.41 ppm for ^1^H**α** +3.1 for ^13^C**α**) to 100% -strand (+0.41 for ^1^H**α**, −3.1 for ^13^C**α**). N- and C-terminal residues, which are perturbed by end effects, are not plotted. **C**. The ^3^J_HNHA_ coupling constants for non-terminal, non-glycine residues are plotted as black diamonds. Dotted lines represent values for **-strand**, **polyproline II helix** and **-helix** (Pardi *et al.*, 1984).

NOE correlations are observed between Tyr and Arg, and between Arg and Trp sidechain nuclei (**Figure 6**). Although these were also observed in RGG3FUS, the designed RGGmini sequence affords site-specific assignments. These signals indicate structural contacts within 0.5 nm which are compatible with a polyproline II helical structure if the side chains are spaced *i, i+3* in sequence. To further assess these contacts, 2D ^13^C-^15^N CON and 1D ^1^H NMR spectra were recorded over temperatures ranging from 5 to 60 °C at pH 10, and over pH values ranging from (2.4 – 9.9) at 5 °C. The CON spectra, which monitor the ^13^CO-^15^N correlations of Gly2-Gly3 and Gly8-Gly9, show essentially no changes upon varying the pH (data not shown) and small linear chemical shift changes as a function of temperature (**Sup Fig. 3A**). These observations are different from those reported by Lim and Hsu (2003) for the peptide YGRKKRRQRRRP, which adopts an isolated polyproline II helix at low temperatures. The backbone ^15^N of that peptide showed significant chemical shifts with respect to coil values which decreased as the temperature is raised. The difference between the results may be due to the low backbone solvent accessibility of the YGRKKRRQRRRP peptide, dominated by residues with bulky side chains versus the relatively exposed backbone of the glycine-rich RGGmini peptide examined here.

**Figure 6:**
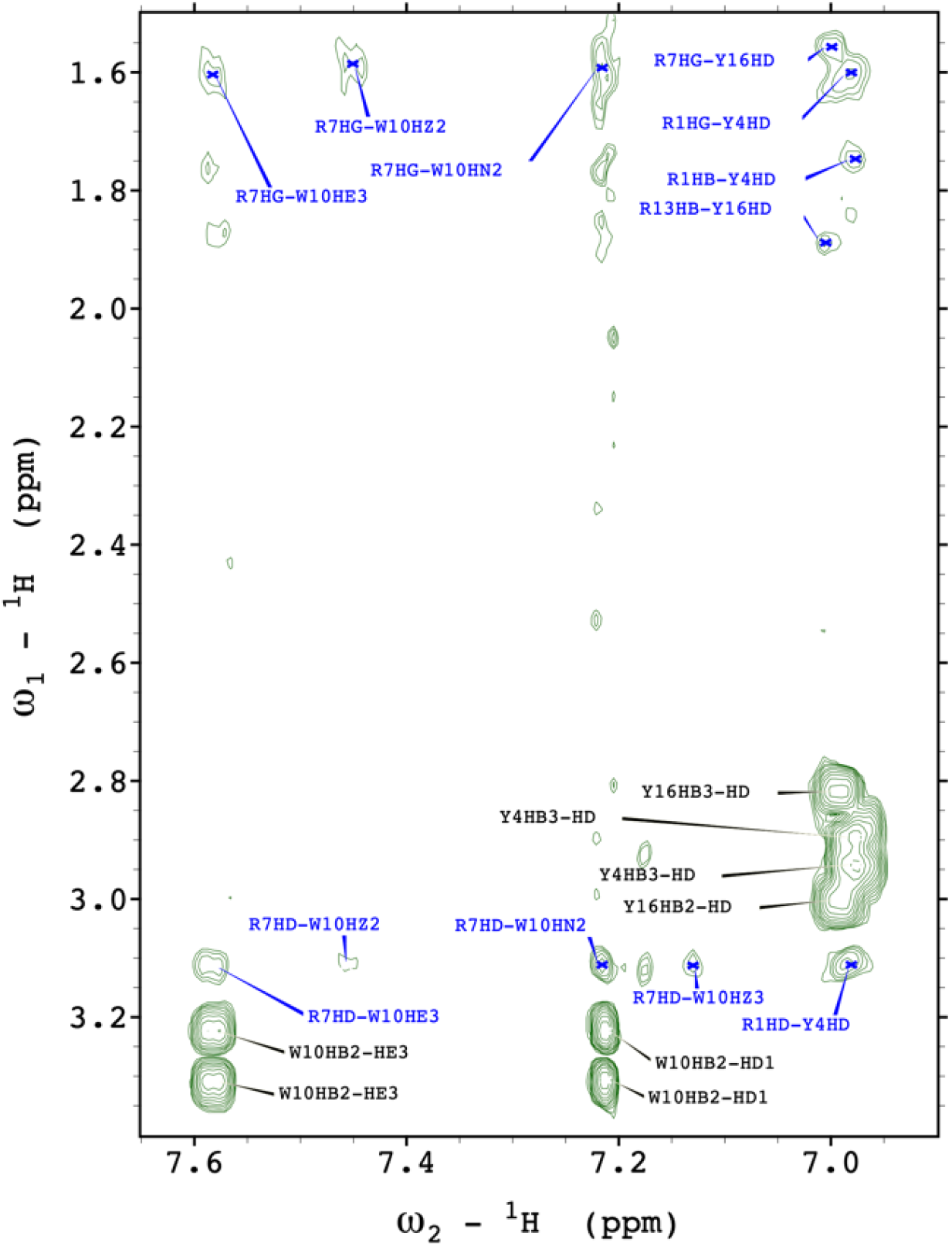
NMR Characterization of RGGmini (II) 2D ^1^H-^1^H NOESY spectrum recorded at pH 9.95 on 6.1 mM RGGmini using a 200 ms mixing time, showing the spectral region of the aromatic to aliphatic correlations. Multiplication factor for contours = (2)^1/2^, so every two steps is a factor of two. Intra- and inter-residual crosspeaks are labeled in **black** and **blue**, respectively.

In contrast to the backbone, substantial changes are seen for the Tyr and Trp side chain aromatic ^1^H chemical shifts (**Sup. Fig. 3B**), as well as Tyr ^1^Hβ, Trp ^1^Hβ and Arg ^1^Hδ and to a minor degree Arg ^1^Hγ upon heating (**Sup. Fig. 3C, D**), relative to the reference DSS peak, which was set to 0.000 ppm at each temperature (**Sup. Fig. 3E**). These results can be interpreted as arising from the structural contacts that are present between the Tyr, Arg and Trp side chains at low temperature which break down upon heating due to thermal denaturation.

Modeling shows that structural contacts among the Tyr, Arg and Trp side chains would be suboptimal in an α-helix (**Figure 7A**) and non-existent with β-strand conformations (**Figure 7B)**. In contrast, these side chains would be close in space in a polyproline II helix, as they are placed three apart in the peptide sequence. Using these NOE signals to derive distance restraints, and ϕ,ψ dihedral angles restrictions derived from the polyproline II helical conformation measured by CD, FTIR, conformational chemical shifts and ^3^J_HNHA_ values, structural models for RGGmini were calculated using CYANA, and are shown in **Figure 7C** and **Supporting Movie 1**. The structural constraints used as input for the calculation and a summary of the output are presented in **Tables S6** and **S7**, respectively. The models show no consistent restraint violations, low CYANA target function energies, and all the **ϕ, ψ** angles are in the most favored regions of the Ramachandran plot. The resulting model structures show a curved polyproline II helix and contacts among the Tyr, Arg and Trp residues. The backbone bending appears to improve contacts between the aromatic and cationic side chains.

**Figure 7:**
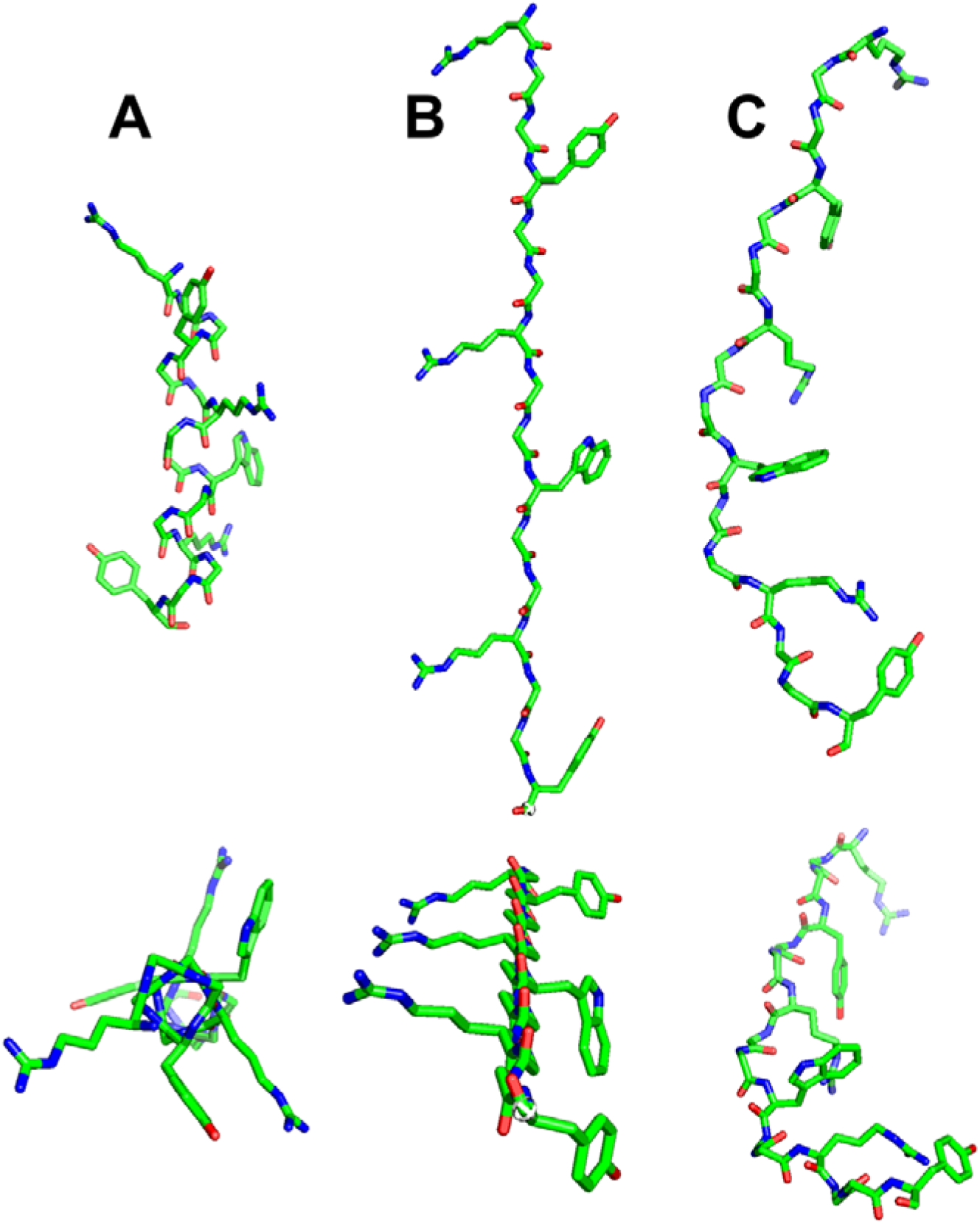
Conformers of RGGmini. **A**. The RGGYGGRGGWGGRGGY peptide modeled as an **α**-helix. **B**. The RGGYGGRGGWGGRGGY peptide modeled as a **β**-strand. The conformers shown in **A** and **B** are incompatible with the experimental data **C**. A representative conformer calculated on the basis of spectral data, which shows a polyproline II helix featuring contacts among the Tyr, Arg and Trp residues. Note that in addition to polyproline II conformers, the structural ensemble includes disordered conformers. Two views of each conformer, rotated 70 to 90° along the x-axis, are shown in panels **A**, **B** and **C**.

## Discussion

Biomolecular condensates are stabilized by a myriad of weak, fluctuating interactions (Wang *et al.*, 2018). Some of these, like hydrogen bonds, cation-π or π-π interactions, form between a few atoms. By contrast, others such as β-strand pairing and the formation of coiled-coils by α-helices arise from elements of secondary structure. Previous investigations have demonstrated that polyproline II helices can also contribute to biomolecular condensate formation by binding to SH3 domains (Li *et al.*, 2012 & Amaya *et al.* 2018). Recently, we determined the NMR spectral parameters of polyproline II helical bundles and predicted the presence of polyproline II helices in the glycine-rich low-complexity regions of FUS and nucleolin (Treviño *et al.*, 2018).

The results reported here show that these sequences, exemplified here by the peptide AcRGGYGGRGGWGGRGGYNH_2_, can adopt polyproline II helices which are stabilized by cation-π interactions amid an ensemble which will also contain disordered conformations. No significant populations of α-helix or β-strands are detected. The backbone of this peptide adopts a polyproline II helix at both low and high pH, as shown by CD or FTIR spectroscopies.

The FUS third RGG region has been shown to make important contributions to FUS liquid/liquid phase separation (Wang *et al.*, 2018). Here, we show that it may transiently adopt isolated single turns of polyproline II helix stabilized b*y i, i+3* interactions and spaced by flexible elbows. Whereas the population of these conformers is low *in vitro*, the *in vivo* stability of the polyproline II conformation could be augmented and contribute to biomolecular condensate formation by alternative interactions, such as interactions with RNA, whose negatively charged phosphate groups are positioned to interact favorably with the positively charged Arg residues of third RGG region. Indeed, RGG regions are known to associate with RNA, including quadruplex structures, through cation-anion and π-stacking interactions (Chong *et al.*, 2018). The FUS protein sequence contains two other RGG stretches called RGG1 and RGG2. RGG1 spans residues 222 – 269 and residues R_259_GGFNKFGGP_268_ could adopt a polyproline II helix stabilized by *i, i+3* side chain interactions, based on the results reported here for RGG3. RGG2, which is composed of residues 371-418, and includes tracts of residues P_391_MGRGGY_397_ and R_396_GGFP_400_, may similarly form polyproline II helices. In addition, RGG2 contains a stretch: R_371_RADFNR_377_, predicted by a helical wheel projection to form a polyproline II helix stabilized by multiple sidechain interactions. In a recent NMR study, this stretch was shown to enhance the binding of the RRM domain of FUS to RNA and to have ^13^Cα, ^13^Cβ and ^1^Ht chemical shifts interpreted as indicative of disorder (Loughlin *et al.*, 2019), but which would also be consistent with a polyproline II helix. Finally, if the G/S,Y,G/S repeats of the N-terminal region of FUS were to also adopt a polyproline II helix, their Tyr residues would be ideally positioned to interact with Arg residues in RGG polyproline II helices. Such associations might help explain why interaction-prone or “sticker” residues tend to be evenly distributed throughout low complexity sequences (Martin *et al.*, 2020). On the other hand, the disordered N-terminal region of Dead box protein 4 which forms germ granules (Nott *et al.*, 2015), while rich in Arg, Gly and aromatic residues, generally lacks the spacings expected to favor polyproline II helix formation.

In addition to these mechanisms, the association of two or more polyproline II helices to form helical bundles, which may be stabilized by the formation of conventional N-H**⋯**O=C as well as weak Cα-H⋯O=C hydrogen bonds, could also contribute to biomolecular condensate formation *in vivo* (Treviño *et al.*, 2018). Finally, related glycine rich sequences could adopt a polyproline helix and bind to profilin protein or OCRE, WW or SH3 domains. In fact, it has been recently shown that the Fyn SH3 domain binds the C-terminal residues of hnRNPA2 (See **Fig. 4 I** of Amaya *et al.*, 2018). Although this hnRNPA2 segment: S_329_-GGSGGYGGRSRY_341_ lacks proline residues, it could adopt a polyproline II helix similar to that characterized here for the RGGmini peptide derived from FUS.

## Materials and Methods

### Peptide Design and Synthesis

The model peptides RGG3FUS and RGGmini have the sequences: RGGRGGYDRGGYRGGRGGDRGNYRGGRGGD and AcRGGYGGRGGWGGRGGYNH_2_, respectively, where Ac- and -NH_2_ represent N-terminal acetyl and C-terminal amide blocking groups, were prepared by standard fMoc Merrifield solid-phase peptide synthesis using a Focus XC peptide synthesizer at the Protein Chemistry Laboratory of the “Margarita Salas” Centre for Biological Investigation of the Spanish National Research Council. The RGGmini peptide labeled with ^13^CO at Gly2 and Gly8 and ^15^N at Gly3 and Gly9 was prepared in the same fashion by substituting isotopically labeled Fmoc amino acids, purchased from Cambridge Isotope Labs, at these positions.

### Nuclear Magnetic Resonance (NMR) Spectroscopy

All spectra were recorded at 5.0 °C on a Bruker 600 MHz (^1^H) Avance spectrometer, unless noted below. The peptide RGG3FUS was characterized by three sets of 2D ^1^H-^1^H COSY, TOCSY and NOESY spectra at pH 2.53, 7.25 and 9.96, recorded in 90% H_2_O / 10% D_2_O. The peptide concentration was 3.1 mM. The initial pH of the sample was 2.53 and the pH was increased by adding 0.1 M sodium carbonate. These spectra were analyzed along the lines described by Wüthrich (1983) to obtain resonance assignments and ^1^HN-^1^Ht coupling constants. Finally, a 2D ^1^H-^13^C HSQC spectrum (natural abundance ^13^C) was recorded at pH 9.96 in D_2_O. The spectral parameters are listed in **Table 2**.

**Table 2:** NMR spectral parameters

| Spectrum | Dimension | Number of scans | Sweep width (in ppm) | Mixing time (in ms) |
| --- | --- | --- | --- | --- |
| 1D $^1\text{H}$ | 32k | 48 | 12 | -- |
| 2D $^1\text{H}$-$^1\text{H}$ COSY | 2048 x 512 | 8 | 11 x 11 | -- |
| 2D $^1\text{H}$-$^1\text{H}$ TOCSY | 2048 x 512 | 8 | 11 x 11 | 60 |
| 2D $^1\text{H}$-$^1\text{H}$ NOESY | 2048 x 512 | 80 | 11 x 11 | 80 & 150 |
| 2D $^1\text{H}$-$^{13}\text{C}$ HSQC | 2048 x 128 | 200 | 11 $^1\text{H}$ x 65 $^{13}\text{C}$<br>(aliphatic, center = 46 ppm) | -- |
| 2D $^1\text{H}$-$^{13}\text{C}$ HSQC | 1k x 128 | 64 | 11 $^1\text{H}$ x 35 $^{13}\text{C}$<br>(aromatic, center = 117.5 ppm) | -- |
| 2D $^1\text{H}$-$^{15}\text{N}$ HSQC | 2048 x 64 | 2 | 10 $^1\text{H}$ x 2 $^{15}\text{N}$ | -- |
| 2D "$^{13}\text{C}$,$^{15}\text{N}$"<br>C_con_iasq | 1024 x 48 | 16 | 4 $^{15}\text{N}$ x 4 $^{13}\text{C}$ | Recorded at 5, 15, 25, 35, 45, 60 & 20 °C |
| 3D HNCO | 2024 x 16 x 16 | 4 | 10 $^1\text{H}$ x 4 $^{15}\text{N}$ x 4 $^{13}\text{C}$ | -- |

Regarding RGGmini, an initial series of 1D ^1^H and 2D ^1^H-^1^H COSY, TOCSY and two NOESY (mixing times = 150 ms & 200 ms) spectra, were recorded on 6.1 mM RGGmini in 90% H_2_O / 10% D_2_O, pH 2.7 at 5 °C. The obtained assignments were further corroborated by analysis of a 2D ^1^H-^15^N HSQC spectrum recorded at natural abundance of ^15^N on a Bruker Avance 800 MHz (^1^H) spectrometer at pH 2.7, as well as two 2D ^1^H-^13^C HSQC spectra (natural abundance ^13^C) centered either on the alphatic or aromatic spectral regions, were recorded at 5 °C Additional COSY, TOCSY and two NOESY spectra (mixing times = 150 & 200 ms) and 2D ^1^H-^13^C HSQC spectra (natural abundance ^13^C), were recorded at 5 °C after adjusting the pH to 9.95 with 0.1 M sodium carbonate.

The ^15^N chemical shift has been reported to show significant changes from coil values in an isolated polyproline II helix (Lam & Hsu, 2003). To assess possible changes in the ^15^N chemical shift under basic conditions, where the ^1^HN signal is lost due to solvent exchange, a RGGmini peptide labeled with ^13^C on the carbonyl of Gly 2 and Gly 8 and with ^15^N at Gly 3 and Gly 9 was characterized. First, 2D ^1^H-^15^N HSQC and 3D HNCO spectra were recorded to assign the ^15^N and ^13^C resonances at low pH. Then, a series of ^13^C’-^15^N (CON) spectra based on ^13^C excitation and detection, Bruker pulse sequence: c_CON_IASQ, were recorded at increasing pH values; namely 2.4, 4.2, 5.4, 7.1 and 9.9 at 5 °C. Finally, a series of 1D ^1^H and CON spectra were recorded at 5, 15, 25, 35, 45, 60 and then 20 °C. The spectral parameters utilized are listed in **Table 2**.

NMR spectra were acquired and processed using TOPSPIN 4.0 (Bruker Biospin). The programs NMRFAM-Sparky (Lee *et al.*, 2015) and CYANA 3.98 (Günter & Buchner, 2015) were used to facilitate manual spectral assignment and to calculate structures, respectively. Predicted chemical shift values for statistical coil ensembles were calculated using the parameters tabulated by Kjaergaard and Poulsen (2011) and Kjaergaard *et al.*, (2011), and implimented on the https://spin.niddk.nih.gov/bax/nmrserver/Poulsen_rc_CS/ server at the Bax laboratory. These values were used to calculate conformational chemical shifts.

For RGG3FUS, distance constraints of 7 Å were set between Tyr QE and Arg QD, and Arg QD and Asp OD1 for all residues spaced *i, i+1* or *i, i+3* along the sequence. Moreover, torsional angles were set to −85° to −65° for ϕ and +135° to +165° for ψ for residues which could be stabilized by *i, i+3* sidechain interactions; namely: 4-7, 9-12, 15-18, 19-22 and 31-34. In the case of RGGmini, initial structures obtained using angular restrictions confined between −80° and −70° for ϕ and +140° and +160° for ψ resulted in six constraint violations for residues 9–12, whose backbone tends to bend. Therefore, the calculations were repeated relaxing the constraint by 10°; using −90° to −60° for ϕ and +130° to +170° for ψ for the residues in question.

### Molecular Dynamics Simulations

Three independent simulations were performed with the amber99sb-ildn force field parameters (Lindorff-Larsen *et al.*, 2010) using GROMACS (Hess *et al.*, 2008). The lowest-energy conformer calculated by CYANA corresponding to RGG3FUS was placed in cubic box filled with TIP3P water (Jorgensen *et al.*, 1983), leaving at least 1.2 nm between each protein atom and the box edges. Following steepest descent energy minimization, the system was equilibrated in two consecutive periods, where the protein atoms were restrained to allow solvent molecules to equilibrate. This was accomplished by running for 500 ps and 2 ns under the NVT and NpT ensembles, respectively, using a modified Berendsen thermostat and the Parrinello-Rahman barostat (Berendsen *et al.*, 1984; Parrinello and Rahman 1981). Molecular Dynamics simulations were run for 65-75 ns using a time step of 2 fs, as enabled by the LINCS algorithm (Hess *et al.*, 1997), and under the NpT ensemble controlling the temperature and pressure with the Nosé-Hoover (Nosé 1984) thermostat (time constant of 0.5 ps) and Parrinello–Rahman (Parrinello and Rahman 1981) barostat (time constant of 1.0 ps). Short-range nonbonded interactions were cut-off at 1 nm, while long-range electrostatics were calculated with the particle mesh Ewald (PME) algorithm (Darden *et al.*, 1993). Dispersion-correction was applied to account for van der Waals interactions at distances longer than the cut off, and periodic boundary conditions were applied in all directions. Different initial velocities were used in each run to allow the system to evolve through distinct trajectories, such that the ~215 ns of total sampling correspond to the sum of independent simulations. The choice of the amber99sb-ildn force field was based on recent literature showing that its parameters do not result in oversampling of PPII conformations (Ouying *et al.*, 2018; Uluca *et al.*, 2018).

### UV Circular Dichroism (CD) Spectroscopy

CD spectra were acquired using a Jasco 810 spectropolarimeter fitted with a Peltier temperature control unit. Far UV-CD spectra were recorded using a 1.0 nm bandwidth scanning from 260 to 210 nm at 20 nm·min^−1^. Two scans were averaged per spectrum. Spectra were recorded at pH 3, 8, and 10, in a 1 mm pathlength cuvette with a peptide concentration of 1.0 mM at 0 °C, 30 °C and 60 °C and finally a repeat spectrum was recorded at 0 °C after recooling. Far-UV CD spectra were recorded at pH 3 and pH 8, where Tyr residues are in the neutral phenol state. Near UV-CD spectra were recorded on a 3.1 mM sample of RGGmini in a 10 mm pathlength cuvette at pH 10 scanning from 320 to 260 nm at a 20 nmomin^−1^ scan speed and a 1.2 nm bandwidth.

### Fourier Transform Infrared (FTIR) Spectroscopy

A Bruker Verex 80 spectrometer equipped with a Specac variable temperature holder and ZnSe windows was used to record FTIR spectra. The sample concentration was 15 mg·mL^−1^, which is equivalent to 9.6 mM. Spectra were recorded in triplicate using 256 scans under four sets of conditions; namely: 5°C, pH 3.5; 5°C, pH 10; 60°C, pH 3.5; and 60°C, pH 10.

## Acknowledgments

This study was supported by projects: LCF/BQ/PR19/11700003 from “La Caixa Foundation” (ID 100010434) to M.M., BBSRC CASE award with UCB Pharma BB/L014734/1 to AJD, and SAF2016-76678-C2-2-R, 2019AEP121 and BTC-PID2019-109306RB-I00 to DVL from the Spanish Ministry of Economy and Competitivity, the Spanish National Research Council and the Spanish Ministry of Science and Innovation, respectively. The authors acknowledge FCT-Portugal for the PhD studentship attributed to Sara S. Félix (PD/BD/148028/2019). NMR experiments were performed in the “Manuel Rico” NMR Laboratory (LMR) of the Spanish National Research Council (CSIC), a node of the Spanish Large-Scale National Facility (ICTS R-LRB). We are grateful to Dr. José Varela Espinosa, Emilia Aporta Sosa and Cristina Quevedo Sierra (MS-CIB / CSIC) for expert technical work with peptide synthesis.

## Supporting Information

### Supporting Figures & Tables

**Figure S1:**
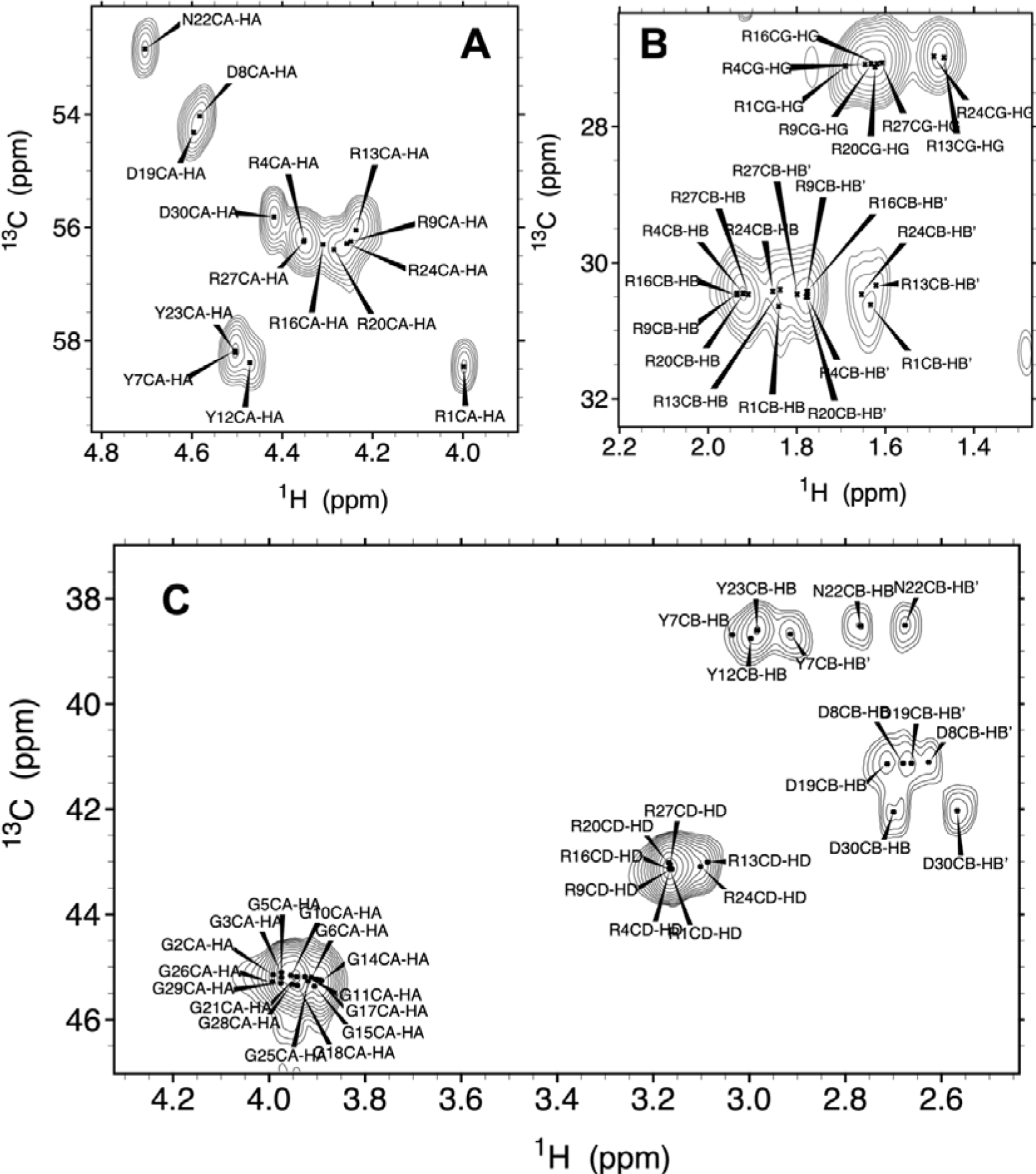
2D ^1^H-^13^C HSQC Spectrum of RGG3FUS at pH 9.96, 5°C in D_2_O. **A.** Downfield ^1^Hα-^13^Cα region. **B.** Upfield region. **C.** Midfield region.

**Figure S2:**
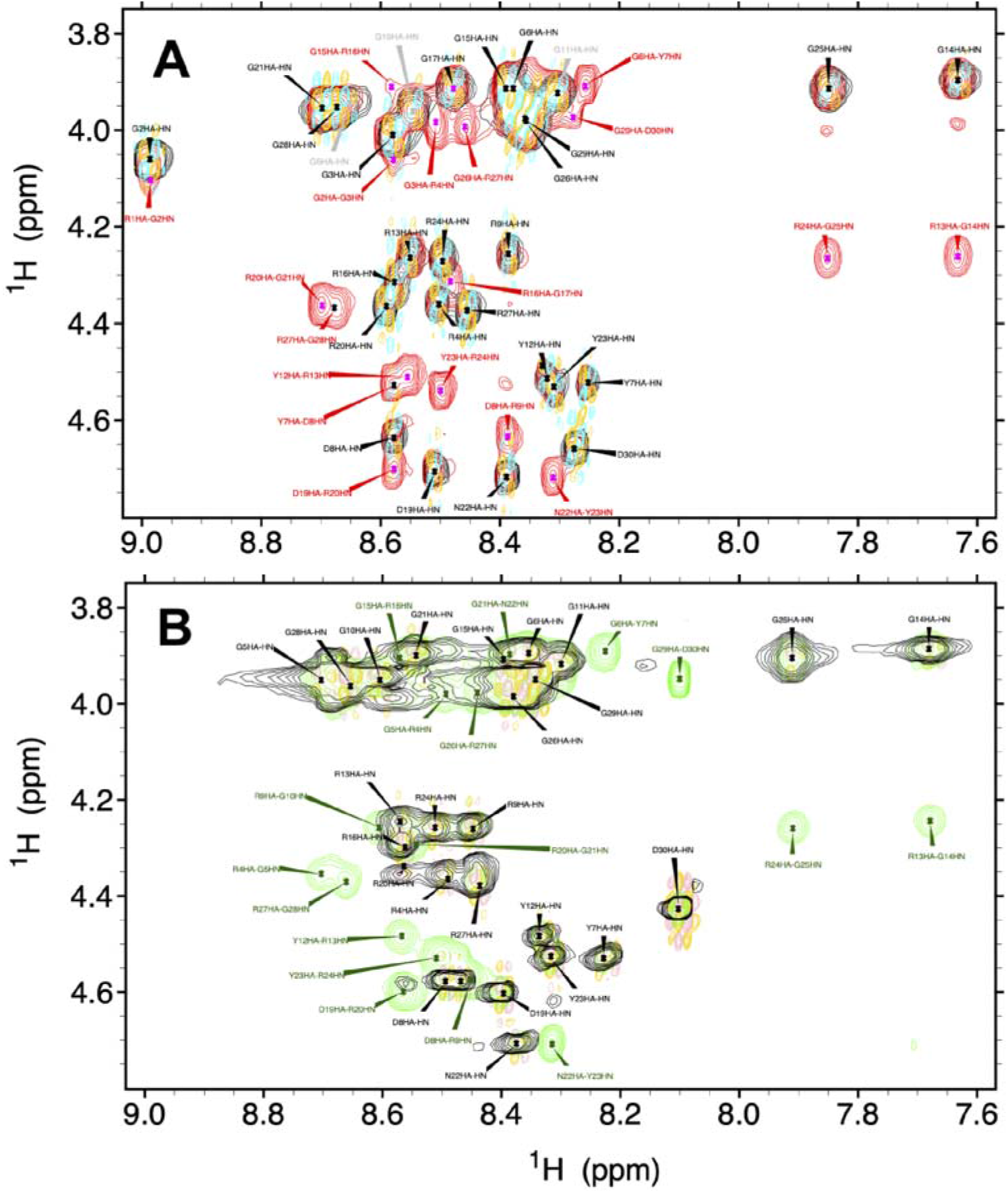
^1^HN-^1^Hα Zone of Spectra of RGG3FUS at pH 2.53 and 7.25, 5°C in 90% H_2_O, 10% D2_O_. **A.** pH 2.53 COSY (+ **gold**, -**cyan**), TOSCY (**black**), NOESY (**red**). Intraresidual assignments are labeled in **black** and interresidue assignments are colored **red**. Tentative assignments are colored **gray**. **B.** pH 7.25 COSY (+ **gold**, -**pink**), TOSCY (**black**), NOESY (**green**). Intraresidual assignments are labeled in **black** and interresidue assignments are colored **green**.

**Figure S3:**
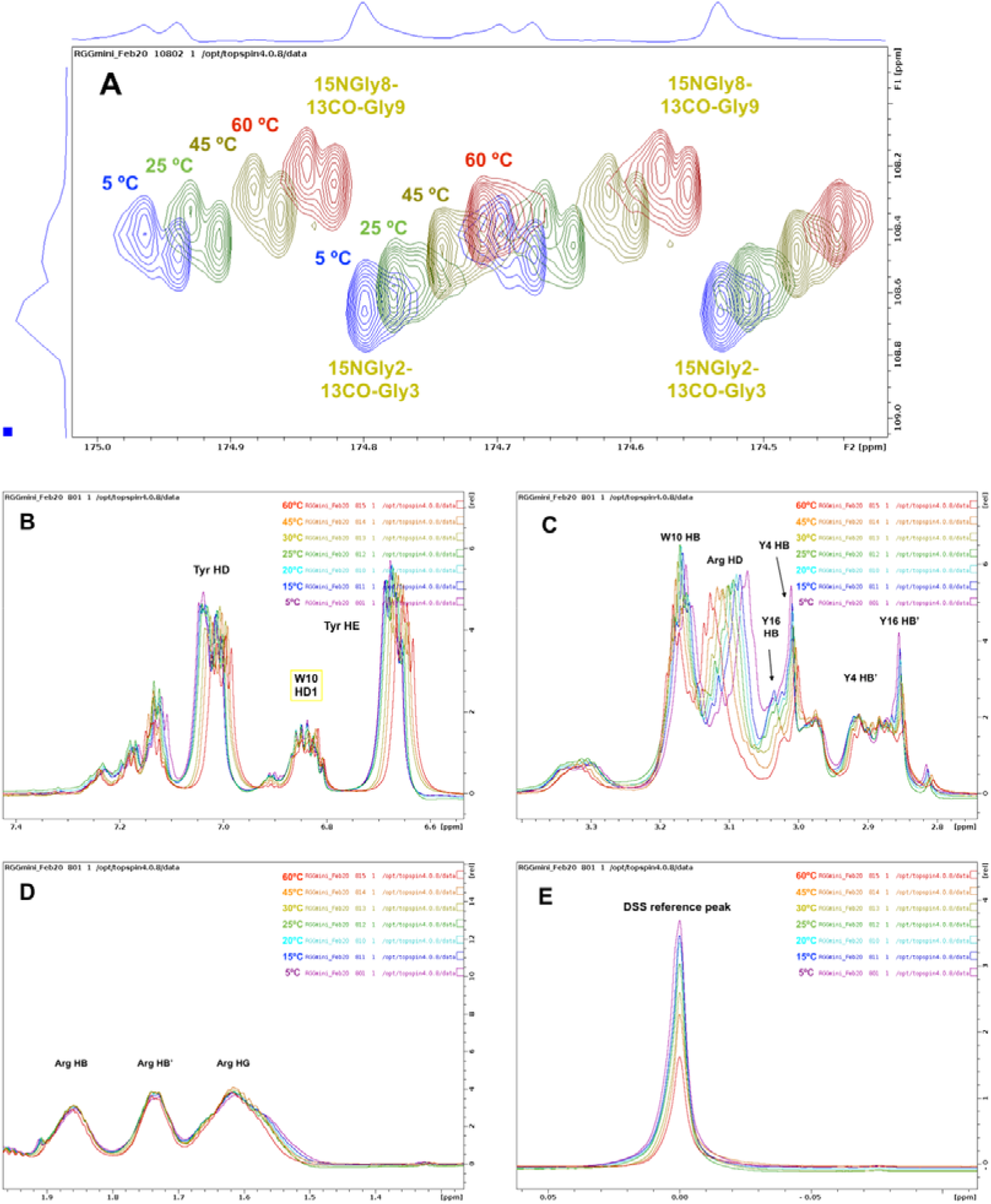
NMR spectra at pH 10 and temperatures ranging from 5 to 60 °C. **A.** 2D CON spectra of RGGYGGRGGWGGRGGY labeled with ^13^C0 at G2 and G8 and ^15^N at G3 and G9. Small, linear change in ^15^N and ^13^C0 chemical shifts with temperature can be detected due to the high resolution of these spectra. The doubled nature of the peaks is due to spin coupling. B. Changes in aromatic ^1^H detected by 1D ^1^H spectra. **C.** Changes in Trp, Tyr ^1^Hβ and Arg ^1^Hδ. **D.** Changes in Arg ^1^Hβ and Aig ^1^Hy. E. The trimethyl signal ^1^H of DSS was set to 0.000 ppm and used to indirectly reference the ^13^C and ^15^N chemical shifts at each temperature. In all the panels, the spectra are colored: 5°C= **purple, 15°C=blue, 20°C=, 25°C=green, 30°C= rmbar, 45°C=,60°C=red.**

**Table S1:** NMR Assignments of RGG3FUS at pH 2.53, 5°C^†^

| Residue | <sup>1</sup>H<sub>N</sub> (ppm) | <sup>1</sup>H<sub>α</sub> (ppm) | <sup>1</sup>H<sub>β</sub>(ppm) | <sup>1</sup>H<sub>γ</sub> (ppm) | <sup>1</sup>H<sub>δ</sub> (ppm) | <sup>1</sup>H<sub>ε</sub> (ppm) |
| --- | --- | --- | --- | --- | --- | --- |
| Arg 1 |  | 4.10 | 1.92, 1.70 | 1.69 | 3.23 | 7.29 |
| Gly 2 | 8.99 | 4.06 |  |  |  |  |
| Gly 3 | 8.51 | 4.00 |  |  |  |  |
| Arg 4 | 8.50 | 4.36 | 1.89, 1.77 | 1.64 | 3.18 | 7.25 |
| Gly 5 | 8.65 | 3.94 |  |  |  |  |
| Gly 6 | 8.38 | 3.91 |  |  |  |  |
| Tyr 7 | 8.26 | 4.52 | 2.99 |  | 7.10 | 6.81 |
| Asp 8 | 8.58 | 4.64 | 2.86, 2.70 |  |  |  |
| Arg 9 | 8.39 | 4.26 | 1.91, 1.77 | 1.64 | 3.18 | 7.25 |
| Gly 10 | 8.55 | 3.96 |  |  |  |  |
| Gly 11 | 8.30 | 3.92 |  |  |  |  |
| Tyr 12 | 8.32 | 4.51 | 2.95 |  | 7.10 | 6.81 |
| Arg 13 | 8.55 | 4.25 | 1.72, 1.64 | 1.51 | 3.11 | 7.19 |
| Gly 14 | 7.64 | 3.90 |  |  |  |  |
| Gly 15 | 8.37 | 3.91 |  |  |  |  |
| Arg 16 | 8.58 | 4.32 | 1.91, 1.78 | 1.64 | 3.18 | 7.25 |
| Gly 17 | 8.48 | 3.91 |  |  |  |  |
| Gly 18 | n.a.<sup>‡</sup> | n.a. |  |  |  |  |
| Asp 19 | 8.51 | 4.70 | 2.90, 2.82 |  |  |  |
| Arg 20 | 8.59 | 4.36 | 1.91, 1.78 | 1.66 | 3.18 | 7.25 |
| Gly 21 | 8.70 | 3.95 |  |  |  |  |
| Asn 22 | 8.39 | 4.71 | 2.79, 2.69 |  | 7.67, 7.01 |  |
| Tyr 23 | 8.31 | 4.54 | 3.01 |  | 7.11 | 6.81 |
| Arg 24 | 8.50 | 4.26 | 1.79, 1.65 | 1.51 | 3.15 | 7.22 |
| Gly 25 | 7.85 | 3.91 |  |  |  |  |
| Gly 26 | 8.36 | 3.99 |  |  |  |  |
| Arg 27 | 8.46 | 4.37 | 1.91, 1.78 | 1.64 | 3.18 | 7.25 |
| Gly 28 | 8.67 | 3.95 |  |  |  |  |
| Gly 29 | 8.35 | 3.97 |  |  |  |  |
| Asp 30 | 8.28 | 4.66 | 2.89 |  |  |  |

<sup>†</sup>In ppm, referencing using the trimethyl singlet of DSS as an internal reference for <sup>1</sup>H.
<sup>‡</sup>n.a.: not assigned

**Table S2:** NMR Assignments of RGG3FUS at pH 7.25, 5°C^†^

| Residue | <sup>1</sup>HN (ppm) | <sup>1</sup>Hα (ppm) | <sup>1</sup>Hβ(ppm) | <sup>1</sup>Hγ (ppm) | <sup>1</sup>Hδ (ppm) | <sup>1</sup>Hε (ppm) |
| --- | --- | --- | --- | --- | --- | --- |
| Arg 1 | n. a.<sup>‡</sup> | 4.00 | 1.84, 1.64 | 1.68 | 3.19 | 7.25 |
| Gly 2 | n. a. | n. a. |  |  |  |  |
| Gly 3 | n. a. | n. a. |  |  |  |  |
| Arg 4 | 8.49 | 4.36 | 1.91, 1.77 | 1.63 | 3.18 | 7.29 |
| Gly 5 | 8.70 | 3.97 |  |  |  |  |
| Gly 6 | 8.35 | 3.89 |  |  |  |  |
| Tyr 7 | 8.22 | 4.53 | 3.04, 2.02 |  | 7.08 | 6.80 |
| Asp 8 | 8.49 | 4.58 | 2.69, 2.62 |  |  |  |
| Arg 9 | 8.45 | 4.26 | 1.91, 1.77 | 1.63 | 3.18 | 7.29 |
| Gly 10 | 8.61 | 3.95 |  |  |  |  |
| Gly 11 | 8.30 | 3.92 |  |  |  |  |
| Tyr 12 | 8.33 | 4.49 | 2.95 |  | 7.10 | 6.80 |
| Arg 13 | 8.57 | 4.24 | 1.74, 1.62 | 1.62 | 3.10 | 7.24 |
| Gly 14 | 7.68 | 3.89 |  |  |  |  |
| Gly 15 | 8.40 | 3.91 |  |  |  |  |
| Arg 16 | 8.57 | 4.30 | 1.93, 1.78 | 1.66 | 3.10 | 7.24 |
| Gly 17 | n.a. | n.a. |  |  |  |  |
| Gly 18 | n.a. | n.a. |  |  |  |  |
| Asp 19 | 8.40 | 4.60 | 2.77, 2.66 |  |  |  |
| Arg 20 | 8.57 | 4.31 | 1.91, 1.77 | 1.63 | 3.18 | 7.29 |
| Gly 21 | 8.54 | 3.90 |  |  |  |  |
| Asn 22 | 8.38 | 4.71 | 2.77, 2.68 |  | 7.70, 7.00 |  |
| Tyr 23 | 8.32 | 4.52 | 3.00 |  | 7.10 | 6.80 |
| Arg 24 | 8.51 | 4.26 | 1.70, 1.67 | 1.51 | 3.14 | 7.25 |
| Gly 25 | 7.91 | 3.91 |  |  |  |  |
| Gly 26 | 8.39 | 3.98 |  |  |  |  |
| Arg 27 | 8.44 | 4.37 | 1.91, 1.77 | 1.63 | 3.18 | 7.29 |
| Gly 28 | 8.66 | 3.96 |  |  |  |  |
| Gly 29 | 8.34 | 3.95 |  |  |  |  |
| Asp 30 | 8.10 | 4.43 | 2.69, 2.57 |  |  |  |

<sup>†</sup>In ppm, referencing using the trimethyl singlet of DSS as an internal reference for <sup>1</sup>H.
<sup>‡</sup> n.a. not assigned

**Table S3:**
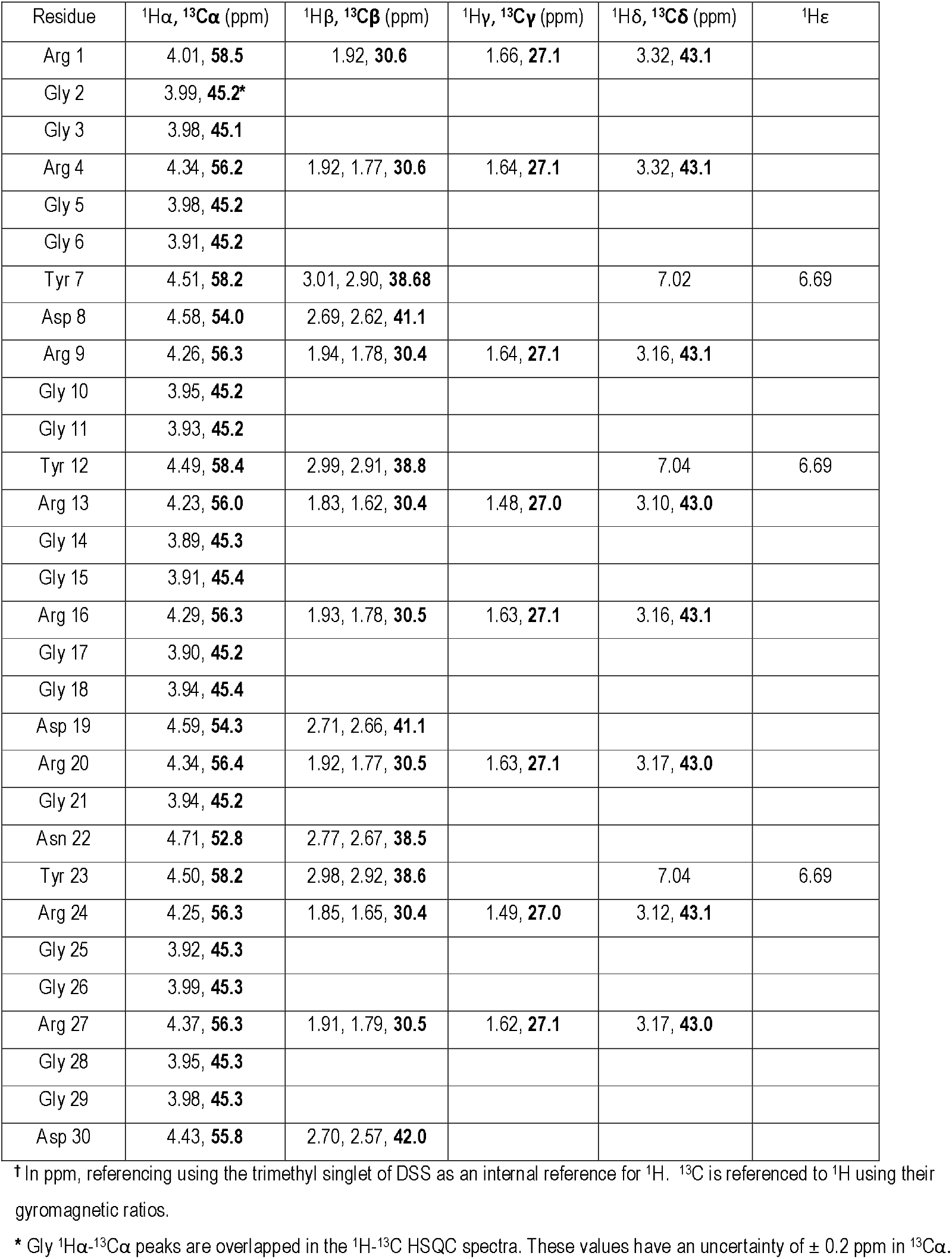
NMR Assignments of RGG3FUS at pH 10, 5°C^†^

| Residue | <sup>1</sup>Hα, <sup>13</sup>Cα (ppm) | <sup>1</sup>Hβ, <sup>13</sup>Cβ (ppm) | <sup>1</sup>Hγ, <sup>13</sup>Cγ (ppm) | <sup>1</sup>Hδ, <sup>13</sup>Cδ (ppm) | <sup>1</sup>Hε |
| --- | --- | --- | --- | --- | --- |
| Arg 1 | 4.01, **58.5** | 1.92, **30.6** | 1.66, **27.1** | 3.32, **43.1** |  |
| Gly 2 | 3.99, **45.2\*** |  |  |  |  |
| Gly 3 | 3.98, **45.1** |  |  |  |  |
| Arg 4 | 4.34, **56.2** | 1.92, 1.77, **30.6** | 1.64, **27.1** | 3.32, **43.1** |  |
| Gly 5 | 3.98, **45.2** |  |  |  |  |
| Gly 6 | 3.91, **45.2** |  |  |  |  |
| Tyr 7 | 4.51, **58.2** | 3.01, 2.90, **38.68** |  | 7.02 | 6.69 |
| Asp 8 | 4.58, **54.0** | 2.69, 2.62, **41.1** |  |  |  |
| Arg 9 | 4.26, **56.3** | 1.94, 1.78, **30.4** | 1.64, **27.1** | 3.16, **43.1** |  |
| Gly 10 | 3.95, **45.2** |  |  |  |  |
| Gly 11 | 3.93, **45.2** |  |  |  |  |
| Tyr 12 | 4.49, **58.4** | 2.99, 2.91, **38.8** |  | 7.04 | 6.69 |
| Arg 13 | 4.23, **56.0** | 1.83, 1.62, **30.4** | 1.48, **27.0** | 3.10, **43.0** |  |
| Gly 14 | 3.89, **45.3** |  |  |  |  |
| Gly 15 | 3.91, **45.4** |  |  |  |  |
| Arg 16 | 4.29, **56.3** | 1.93, 1.78, **30.5** | 1.63, **27.1** | 3.16, **43.1** |  |
| Gly 17 | 3.90, **45.2** |  |  |  |  |
| Gly 18 | 3.94, **45.4** |  |  |  |  |
| Asp 19 | 4.59, **54.3** | 2.71, 2.66, **41.1** |  |  |  |
| Arg 20 | 4.34, **56.4** | 1.92, 1.77, **30.5** | 1.63, **27.1** | 3.17, **43.0** |  |
| Gly 21 | 3.94, **45.2** |  |  |  |  |
| Asn 22 | 4.71, **52.8** | 2.77, 2.67, **38.5** |  |  |  |
| Tyr 23 | 4.50, **58.2** | 2.98, 2.92, **38.6** |  | 7.04 | 6.69 |
| Arg 24 | 4.25, **56.3** | 1.85, 1.65, **30.4** | 1.49, **27.0** | 3.12, **43.1** |  |
| Gly 25 | 3.92, **45.3** |  |  |  |  |
| Gly 26 | 3.99, **45.3** |  |  |  |  |
| Arg 27 | 4.37, **56.3** | 1.91, 1.79, **30.5** | 1.62, **27.1** | 3.17, **43.0** |  |
| Gly 28 | 3.95, **45.3** |  |  |  |  |
| Gly 29 | 3.98, **45.3** |  |  |  |  |
| Asp 30 | 4.43, **55.8** | 2.70, 2.57, **42.0** |  |  |  |

<sup>†</sup>In ppm, referencing using the trimethyl singlet of DSS as an internal reference for <sup>1</sup>H. <sup>13</sup>C is referenced to <sup>1</sup>H using their gyromagnetic ratios.
\* Gly <sup>1</sup>Hα -<sup>13</sup>Cα peaks are overlapped in the <sup>1</sup>H-<sup>13</sup>C HSQC spectra. These values have an uncertainty of $\pm 0.2$ ppm in <sup>13</sup>Cα .

**Table S4:** NMR Assignments of RGGmini at pH 3, 5°C^†^

| Residue | <sup>1</sup>H, <sup>15</sup>N<br>(ppm)<sup>a</sup> | <sup>1</sup>Hα, <sup>13</sup>Cα (ppm) | <sup>1</sup>Hβ, <sup>13</sup>Cβ (ppm) | <sup>1</sup>Hγ, <sup>13</sup>Cγ (ppm) | <sup>1</sup>Hδ, <sup>13</sup>Cδ (ppm) | <sup>1</sup>Hε, <sup>13</sup>Cε,<br><sup>15</sup>Nε (ppm) |
| --- | --- | --- | --- | --- | --- | --- |
| CH<sub>3</sub>-CO-§ |  | 1.99, **25.79**<br>(methyl) |  |  |  |  |
| Arg 1 | | 4.08, **58.25** | 1.92, **30.95** | 1.66, **27.0** ± 0.1 | 3.19, **43.09** | 7.22, **108.3** |
| Gly 2 | 8.90 | 4.01, **45.1\*** |  |  |  |  |
| Gly 3 | 8.44 | 3.87, **45.1\*** |  |  |  |  |
| Tyr 4 | 8.39 | 4.51, **58.37** | 3.01, 2.91, **38.46** |  | 7.07, **133.0** | 6.79, **117.9** |
| Gly 5 | 8.60 | 3.82, **45.1\*** |  |  |  |  |
| Gly 6 | 7.89 | 3.89, **45.1\*** |  |  |  |  |
| Arg 7 | 8.42 | 4.28, 56.27 | 1.87, 1.74, **30.35** | 1.62, 1.56, **27.0**<br>± 0.1 | 3.19, **43.09** | 7.19, **108.3** |
| Gly 8 | 8.62 | 3.91, **45.1\*** |  |  |  |  |
| Gly 9 | 8.25 | 3.87, **45.1\*** |  |  |  |  |
| Trp 10‡ | 8.28 | 4.64, 57.45 | 3.29, 3.21, **29.34** |  |  | 10.17,<br>**129.50** |
| Gly 11 | 8.56, **108.5** | 3.79, **45.1\*** |  |  |  |  |
| Gly 12 | 8.24 | 3.81, **45.1\*** |  |  |  |  |
| Arg 13 | 8.29 | 4.33, **56.09** | 1.90, 1.77, **30.62** | 1.61, **27.0** ± 0.1 | 3.19, **43.09** | 7.19, **108.3** |
| Gly 14 | 8.61 | 3.91, **45.1\*** |  |  |  |  |
| Gly 15 | 7.88 | 3.80, **45.1\*** |  |  |  |  |
| Tyr 16 | 7.94 | 4.45, **59.17** | 3.06, 2.87, **39.21** |  | 7.07, **133.0** | 6.79, **117.9** |
| -NH<sub>2</sub>¶ | 7.75, 7.26,<br>**108.4** |  |  |  |  |  |

<sup>†</sup> In ppm, referencing using the trimethyl singlet of DSS as an internal reference for <sup>1</sup>H. <sup>13</sup>C and <sup>15</sup>N are referenced to <sup>1</sup>H using their gyromagnetic ratios.
§ The N-terminus of the peptide is acetylated. The chemical shifts of the acetyl methyl group come from the <sup>1</sup>H-<sup>13</sup>C HSQC spectrum.
\* Gly <sup>1</sup>Hα -<sup>13</sup>Cα peaks are overlapped in the <sup>1</sup>H-<sup>13</sup>C HSQC spectra. These values have an uncertainty of ± 0.2 ppm in <sup>13</sup>Cα .
‡ The Trp ring assignments are <sup>1</sup>H<sup>13</sup>C $\delta$ 1 = 7.21, **126.8**; <sup>1</sup>H<sup>15</sup>N $\epsilon$ = 10.17, **129.50**; <sup>1</sup>H<sup>13</sup>C $\epsilon$ 3 = 7.58, **120.7**; <sup>1</sup>H<sup>13</sup>C $\zeta$ 3 = 7.13, **121.9**; <sup>1</sup>H<sup>13</sup>C $\eta$ 2 = 7.22, **124.5**; <sup>1</sup>H<sup>13</sup>C $\zeta$ 2 = 7.45, **114.4** ppm.
¶ The C-terminus of the peptide is carboxamidated. The chemical shifts of the terminal -NH<sub>2</sub> moiety come from the <sup>1</sup>H-<sup>15</sup>N HSQC spectrum.

**Table S5:** NMR Assignments of RGGmini at pH 10, 5°C^†^

| Residue | <sup>1</sup>Hα, <sup>13</sup>Cα (ppm) | <sup>1</sup>Hβ, <sup>13</sup>Cβ (ppm) | <sup>1</sup>Hγ, <sup>13</sup>Cγ (ppm) | <sup>1</sup>Hδ, <sup>13</sup>Cδ (ppm) | <sup>1</sup>Hε, <sup>13</sup>Cε,<br><sup>15</sup>Nε (ppm) |
| --- | --- | --- | --- | --- | --- |
| CH<sub>3</sub>-CO-<sup>‡</sup> | 1.99, **25.79**<br>(methyl) |  |  |  |  |
| Arg 1 | 4.08, **58.25** | 1.89, 1.75, **30.95** | 1.60, **27.0** | 3.11, **43.1** |  |
| Gly 2 | 3.93, **45.1\*** |  |  |  |  |
| Gly 3 | 3.93, **45.1** |  |  |  |  |
| Tyr 4 | 4.47, **58.38** | 3.03, 2.92, **38.45** |  | 7.00, **133.2** | 6.68, **119.6** |
| Gly 5 | 3.93, **45.1** |  |  |  |  |
| Gly 6 | 3.93, **45.1** |  |  |  |  |
| Arg 7 | 4.34, **56.27** | 1.95, 1.83, **30.35** | 1.60, **27.0** | 3.11, **43.1** |  |
| Gly 8 | 3.93, **45.1** |  |  |  |  |
| Gly 9 | 3.93, **45.1** |  |  |  |  |
| Trp 10<sup>‡</sup> | 4.72, **57.45** | 3.31, 3.29, **29.34** |  |  |  |
| Gly 11 | 3.93, **45.1** |  |  |  |  |
| Gly 12 | 3.93, **45.1\*** |  |  |  |  |
| Arg 13 | 4.22, **56.09** | 1.97, 1.85, **30.61** | 1.70, **27.0** | 3.11, **43.1** |  |
| Gly 14 | 3.93, **45.** |  |  |  |  |
| Gly 15 | 3.93, **45.1** |  |  |  |  |
| Tyr 16 | 4.42, **59.17** | 3.05, 2.86, **39.22** |  | 7.00, **133.2** | 6.68, **119.6** |

<sup>†</sup> In ppm, referencing using the trimethyl singlet of DSS as an internal reference for <sup>1</sup>H. <sup>13</sup>C and <sup>15</sup>N are referenced to <sup>1</sup>H using their gyromagnetic ratios.
\* Gly <sup>1</sup>Hα -<sup>13</sup>Cα peaks are overlapped in the <sup>1</sup>H-<sup>13</sup>C HSQC spectra. These values have an uncertainty of $\pm 0.4$ ppm in <sup>13</sup>Cα .
<sup>‡</sup> The Trp <sup>1</sup>H, <sup>13</sup>C ring assignments are $\delta 1 = 7.20$ , **127.2**, $\epsilon 3 = 7.56$ , **120.9**, <sup>1</sup>H $\zeta 3 = 7.11$ , **122.1**, <sup>1</sup>H $\eta 2 = 7.20$ , **124.7**, $\zeta 2 = 7.44$ , **114.7**.
<sup>¶</sup> The C-terminus of the peptide is carboxamidated. The chemical shifts of the terminal -NH<sub>2</sub> moiety come from the <sup>1</sup>H-<sup>15</sup>N HSQC spectrum.

**Table S6:**
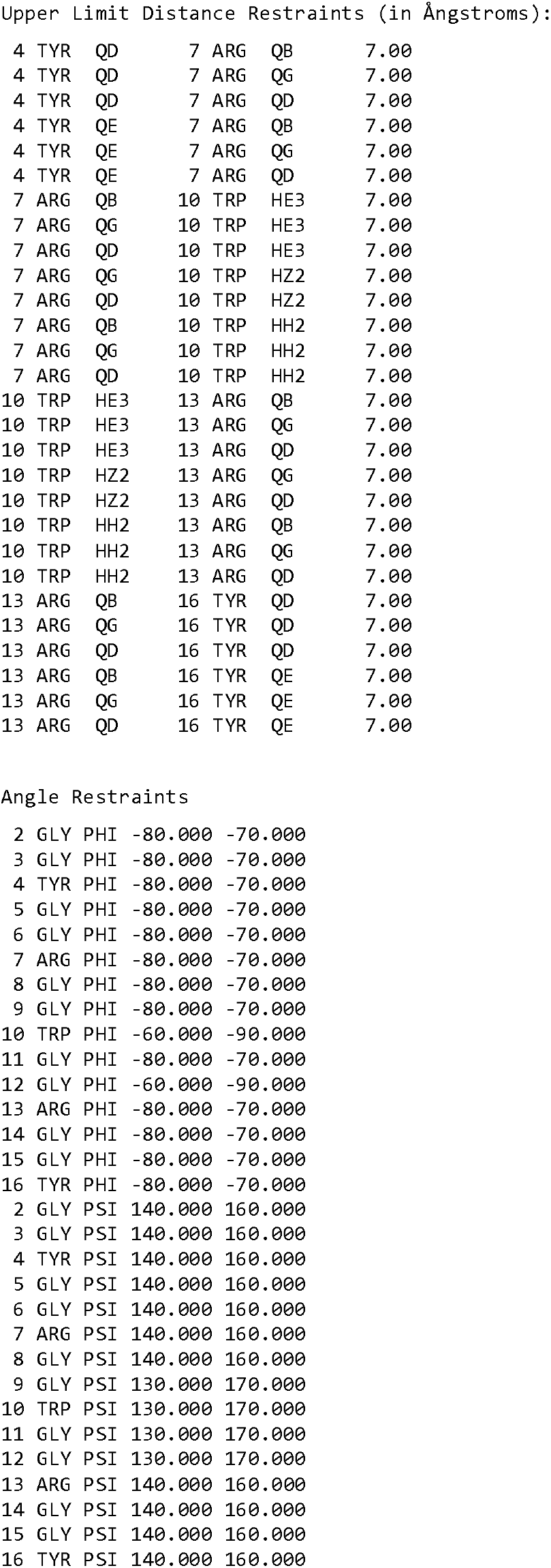
Restraints for Structure Calculations Using CYANA

Upper Limit Distance Restraints (in Ångstroms):
|  |  |  |  |  |  |  |
| --- | --- | --- | --- | --- | --- | --- |
| 4 | TYR | QD | 7 | ARG | QB | 7.00 |
| 4 | TYR | QD | 7 | ARG | QG | 7.00 |
| 4 | TYR | QD | 7 | ARG | QD | 7.00 |
| 4 | TYR | QE | 7 | ARG | QB | 7.00 |
| 4 | TYR | QE | 7 | ARG | QG | 7.00 |
| 4 | TYR | QE | 7 | ARG | QD | 7.00 |
| 7 | ARG | QB | 10 | TRP | HE3 | 7.00 |
| 7 | ARG | QG | 10 | TRP | HE3 | 7.00 |
| 7 | ARG | QD | 10 | TRP | HE3 | 7.00 |
| 7 | ARG | QG | 10 | TRP | HZ2 | 7.00 |
| 7 | ARG | QD | 10 | TRP | HZ2 | 7.00 |
| 7 | ARG | QB | 10 | TRP | HH2 | 7.00 |
| 7 | ARG | QG | 10 | TRP | HH2 | 7.00 |
| 7 | ARG | QD | 10 | TRP | HH2 | 7.00 |
| 10 | TRP | HE3 | 13 | ARG | QB | 7.00 |
| 10 | TRP | HE3 | 13 | ARG | QG | 7.00 |
| 10 | TRP | HE3 | 13 | ARG | QD | 7.00 |
| 10 | TRP | HZ2 | 13 | ARG | QG | 7.00 |
| 10 | TRP | HZ2 | 13 | ARG | QD | 7.00 |
| 10 | TRP | HH2 | 13 | ARG | QB | 7.00 |
| 10 | TRP | HH2 | 13 | ARG | QG | 7.00 |
| 10 | TRP | HH2 | 13 | ARG | QD | 7.00 |
| 13 | ARG | QB | 16 | TYR | QD | 7.00 |
| 13 | ARG | QG | 16 | TYR | QD | 7.00 |
| 13 | ARG | QD | 16 | TYR | QD | 7.00 |
| 13 | ARG | QB | 16 | TYR | QE | 7.00 |
| 13 | ARG | QG | 16 | TYR | QE | 7.00 |
| 13 | ARG | QD | 16 | TYR | QE | 7.00 |

Angle Restraints
|  |  |  |  |  |
| --- | --- | --- | --- | --- |
| 2 | GLY | PHI | -80.000 | -70.000 |
| 3 | GLY | PHI | -80.000 | -70.000 |
| 4 | TYR | PHI | -80.000 | -70.000 |
| 5 | GLY | PHI | -80.000 | -70.000 |
| 6 | GLY | PHI | -80.000 | -70.000 |
| 7 | ARG | PHI | -80.000 | -70.000 |
| 8 | GLY | PHI | -80.000 | -70.000 |
| 9 | GLY | PHI | -80.000 | -70.000 |
| 10 | TRP | PHI | -60.000 | -90.000 |
| 11 | GLY | PHI | -80.000 | -70.000 |
| 12 | GLY | PHI | -60.000 | -90.000 |
| 13 | ARG | PHI | -80.000 | -70.000 |
| 14 | GLY | PHI | -80.000 | -70.000 |
| 15 | GLY | PHI | -80.000 | -70.000 |
| 16 | TYR | PHI | -80.000 | -70.000 |
| 2 | GLY | PSI | 140.000 | 160.000 |
| 3 | GLY | PSI | 140.000 | 160.000 |
| 4 | TYR | PSI | 140.000 | 160.000 |
| 5 | GLY | PSI | 140.000 | 160.000 |
| 6 | GLY | PSI | 140.000 | 160.000 |
| 7 | ARG | PSI | 140.000 | 160.000 |
| 8 | GLY | PSI | 140.000 | 160.000 |
| 9 | GLY | PSI | 130.000 | 170.000 |
| 10 | TRP | PSI | 130.000 | 170.000 |
| 11 | GLY | PSI | 130.000 | 170.000 |
| 12 | GLY | PSI | 130.000 | 170.000 |
| 13 | ARG | PSI | 140.000 | 160.000 |
| 14 | GLY | PSI | 140.000 | 160.000 |
| 15 | GLY | PSI | 140.000 | 160.000 |
| 16 | TYR | PSI | 140.000 | 160.000 |

**Table S7:**
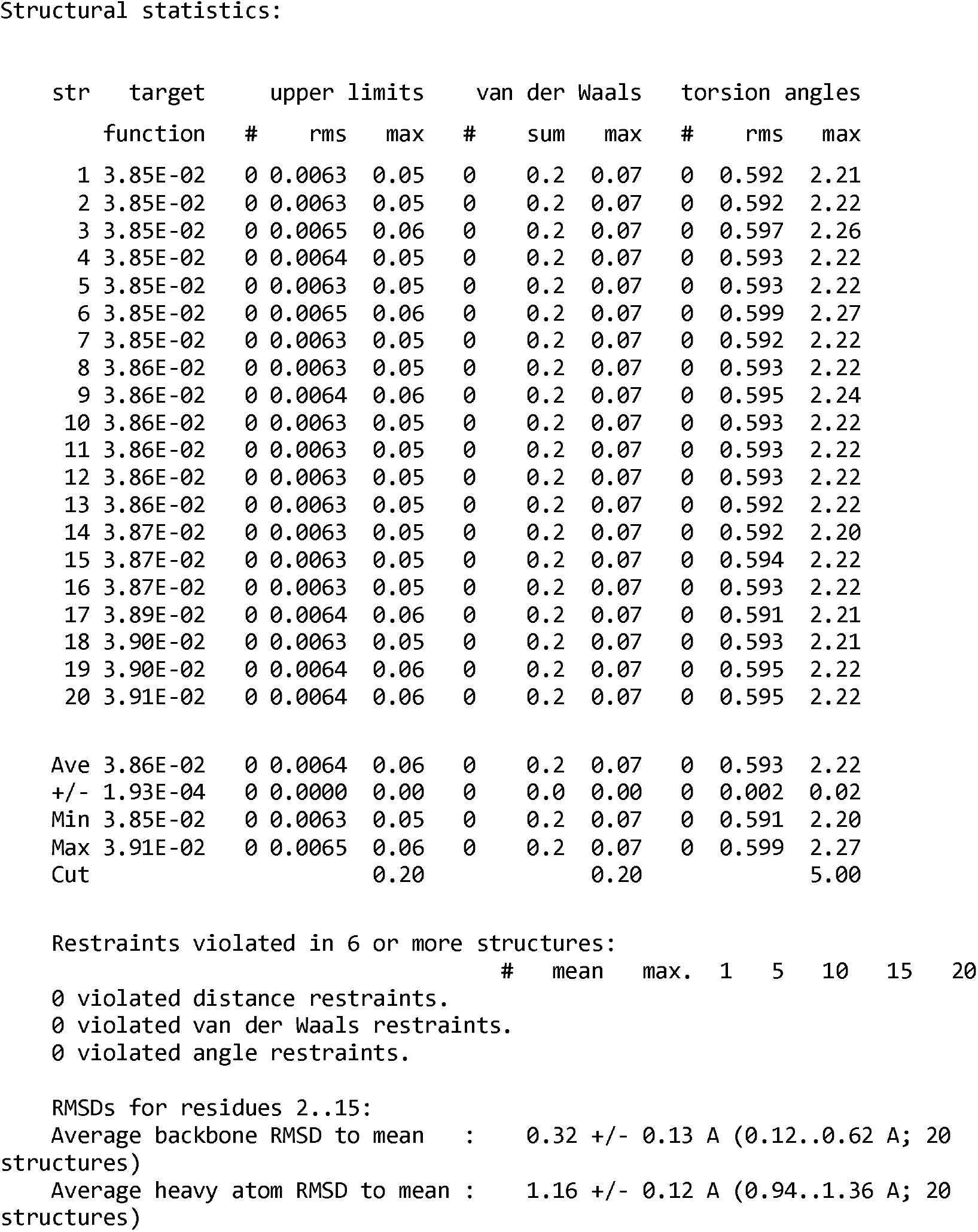
Output Statistics for RGGmini CYANA Structure Calculation

